# Population genomic evidence for a repeated introduction and rapid expansion in Europe of a maize fungal pathogen

**DOI:** 10.1101/2020.09.18.303354

**Authors:** Mireia Vidal-Villarejo, Fabian Freund, Hendrik Hanekamp, Andreas von Tiedemann, Karl Schmid

## Abstract

Modern agricultural practices and the climate change foster the rapid spread of plant pathogens like the maize fungal pathogen *Setosphaeria turcica*, which causes Northern corn leaf blight and expanded into Central Europe since the 1980s. To investigate the rapid expansion of *S. turcica* we sequenced 121 isolates from Europe and Kenya. Population genomic inference revealed a single genetically diverse cluster in Kenya and three clonal lineages with low diversity and one cluster of multiple clonal sublineages in Europe. Phylogenetic dating suggests that all European lineages originated by sexual reproduction outside Europe and subsequently were subsequently introgressed multiple times. In contrast to Kenyan isolates, European isolates did not show sexual recombination despite the presence of both *MAT1-1* and *MAT1-2* mating types. Coalescent analysis of the geographically most widespread European lineage supported a neutral, strongly exponential population growth model over models with natural selection caused by host defence resistance or environmental adaptation. Within clonal lineages, we observed phenotypic variation in virulence to different monogenic resistances that may originate from repeated mutations in virulence genes. Association mapping between genetic clusters did not identify genomic regions associated with pathogen races but uncovered strongly differentiated genomic regions between clonal lineages that harbor putative effector genes. In conclusion, the expansion and population growth of *S. turcica* in Europe was mainly driven by the expansion of maize cultivation area and not by rapid adaptation.

**Significance statement:** The geographic expansion and plant pathogens caused by modern agricultural practices and climate change is a major problem in modern agriculture. We investigated the rapid spread of the maize fungal pathogen Setosphaeria turcica by whole genome sequencing of isolates from Kenya and Europe and demonstrated that the rapid expansion in Central Europe since the 1980s mainly reflects the rapid growth of the maize cultivation area in this region and not a rapid adaptation to resistant maize varieties. Our analyses show that by monitoring whole genome sequence diversity of plant pathogens and their invasion history, agricultural management and breeding strategies can be developed to control the evolution and future spread of plant pathogens.

## Introduction

Modern agricultural practice is characterized by reduced crop rotation, large field sizes of monocultures, high chemical inputs and cultivation of resistant varieties. These factors influence both short-term epidemics and a long-term evolution of resistant pathogen strains that may rapidly expand over large geographic areas (Papaïx et al., 2017). In addition, climate warming favors the spread and adaptation of pathogen species to new environments and geographic regions (Bebber et al., 2013). These factors contribute to rapid crop-pathogen co-evolution, whose understanding is essential to improve management practices and plant breeding to maintain food security in a rapidly changing world (McDonald and Stukenbrock, 2016). Global pathogen monitoring systems for plant pathogens identify the origin and expansion of new pathogen strains (Islam et al., 2016) to support resistance breeding and adaptation of crop management practices. Disease monitoring is greatly facilitated by genome sequencing to characterize pathogen diversity (Hubbard et al., 2015) although a sequence-based prediction of virulence types remains challenging due to a rapid evolution of pathogen genomes (Lamour et al., 2012; Raffaele et al., 2010;Dong et al., 2015;Thordal-Christensen et al., 2018;Frantzeskakis et al., 2019). Sequencing data were used to track the epidemiology and demographic history of pathogens (e.g.,Gladieux et al.,2018;Stam et al., 2019;Latorre et al., 2020) and to reconstruct introductions (Yoshida et al., 2013). However, the relative importance of demographic effects versus selection-driven adaptation to cultivation conditions or plant resistance genes is still little understood. Therefore, a characterization of demography and selection to evaluate the evolutionary potential of pathogen species (McDonald and Linde, 2002) will contribute to developing evolution-informed, durable crop management strategies to avoid rapid breaking of host resistance genes and reduce chemical inputs in plant protection (Burdon et al., 2014).

The hemibiotrophic fungal pathogen *Setosphaeria turcica* (Luttrell) Leonard and Suggs (teleomorph *Exserohilum turcicum*, formerly known as *Helminthosporium turcicum*) is the most important leaf pathogen of maize. It causes Northern corn leaf blight (NCLB), whose symptoms are long, elliptical stripes of necrotic tissues (lesions) on maize leaves, which limits the photosynthetic productivity and causes yield reduction (Galiano-Carneiro and Miedaner, 2017). NCLB is a worldwide disease with a high incidence in the tropics, where it is a major cause of yield loss in maize. The most important methods for controlling the disease are breeding of resistant varieties (Poland et al., 2011) and adapted management practices including fungicide applications. Additional management practices, such as biological control, are being studied (Sartoria et al., 2015;Sartori et al., 2017). *S. turcica* shows asexual and sexual reproduction, which requires mating of two strains with different *MAT1-1* and *MAT1-2* alleles at the *MAT1* mating type locus. Worldwide surveys of genetic diversity of S. turcica showed that sexual reproduction is restricted to regions with a warm climate (Galiano-Carneiro and Miedaner, 2017). Genetic diversity was higher in populations from Mexico in comparison to Kenya, China and Europe suggesting that *S. turcica* originated in Mexico and recently arrived in Europe (Borchardt et al., 1998). NCLB was first reported in Italy in 1876, followed by South-Western France around 1900. Until the 1980s, NCLB was mainly restricted to the warmer regions of Southern Europe and the Balkans, but between 1988 and 1992 the disease crossed the Alps, and in 1995 it was reported in the Upper Rhine Valley in South Germany. Afterwards it rapidly expanded throughout the maize cultivation regions in Northwestern Europe. In response to the expansion of NCLB in Europe, breeders improved commercial varieties by selecting for polygenic, quantitative resistances and by introgression of monogenic, race-specific resistance genes from genetic resources. The four main resistance genes introgressed are *Ht1, Ht2, Ht3 and Htn1* (Welz and Geiger, 2000). Different races of *S. turcica* are defined by their infection ability of a differentiation set of varieties harboring one of the four *Ht* genes. Race monitoring of more than 500 isolates revealed that *S. turcica* races are unequally distributed throughout Europe (Hanekamp, 2016). Such a distribution raises the question whether the rapid expansion reflects a neutral demographic process like a repeated and independent introduction of different strains that were rapidly distributed by seed trade and agricultural practices, or a selection-driven adaptation to resistant host varieties that favored the rapid expansion of novel, virulent pathogen strains throughout Europe.

We investigated both hypotheses by characterizing the genomic diversity of *S. turcica* isolates collected from natural infections of different susceptible maize varieties lacking known *Ht* genes that were cultivated throughout Europe in 2011 and 2012. Using phylogenetic analyses and coalescence models we identify different clonal lineages throughout Central and Western Europe that are distinct from Kenyan isolates used for comparison. The overall genetic diversity of the most widespread European clonal lineage was not shaped by strong selection exerted by host resistance genes, but reflects a neutral, exponential growth.

## Results

### Read mapping and variant discovery

We sequenced a sample of 166 isolates (157.2 GB raw sequence) from 11 different countries (Supplementary Dataset S1) and subsequently removed 37 isolates because of low coverage or a high proportion of reads not mapping to the reference genome. Eight samples were technical replicates of the same isolate to estimate the sequencing error rate. After excluding low quality samples and replicates we analysed a final sample of 121 isolates with an average read coverage of 14.9x and a range from 5.5x to 44.7x coverage. SNP calling with both GATK and samtools-bcftools identified 55,534 SNPs by both methods, of which 23,209 SNPs were retained after filtering (Materials and Methods). SNPs with a maximum of 35% of missing data were imputed by multiple correspondence analysis (MCA). The median number of SNPs differing between the eight technical replicates was 9.5, which corresponds to 99.96% identity between replicates (Supplementary Table S1). To polarize SNPs into ancestral and derived variants we included *Bipolaris sorokiniana and Bipolaris maydis as outgroups* (Ohm et al., 2012;Condon et al., 2013). This data set was expanded by two *Setosphaeria turcica* reference genomes obtained from isolates Et28A and NY001 collected in the United States, which resulted in a total sample of 123 isolates. The data derived from this sample consisted of 4,257 polarized SNPs, corresponding to 18.3% of non-polarized SNP data.

### Presence of different clonal lineages

To determine the genetic relationship of *S. turcica* isolates we clustered the original 121 samples with ADMIXTURE into *K* = 5 clusters (Fig.1B). Five isolates had ancestry coefficients of <70% and were not assigned to clusters. All clusters defined by ADMIXTURE were supported by a rooted Neighbor-Joining tree based on polarized SNPs, a principal component analysis (PCA) and Community Oriented Network Estimation (CONE;Kuismin et al., 2017), see Fig.1A-D. Three of the five ancestral clusters, which we named ‘Big Clonal’ (47 isolates), ‘Small Clonal’ (16 isolates) and ‘French Clonal’ (9 isolates), showed very short internal branches and the two remaining clusters, ‘Diverse’ (17 isolates) and ‘Kenyan’ (27 isolates), showed long internal branches in the phylogenetic tree. The NJ tree, PCA and Neighbor-Net reveal a close relationship of the French Clonal cluster with the Kenyan isolates and a strong differentiation from the other three European clusters (Fig.1A,C,E and Supplementary Fig. S1). All five clusters, however, appear to have arisen by sexual recombination as indicated by reticulate patterns at the base of each clade in the Neighbor-Net (Fig.1E).

**Fig 1.**
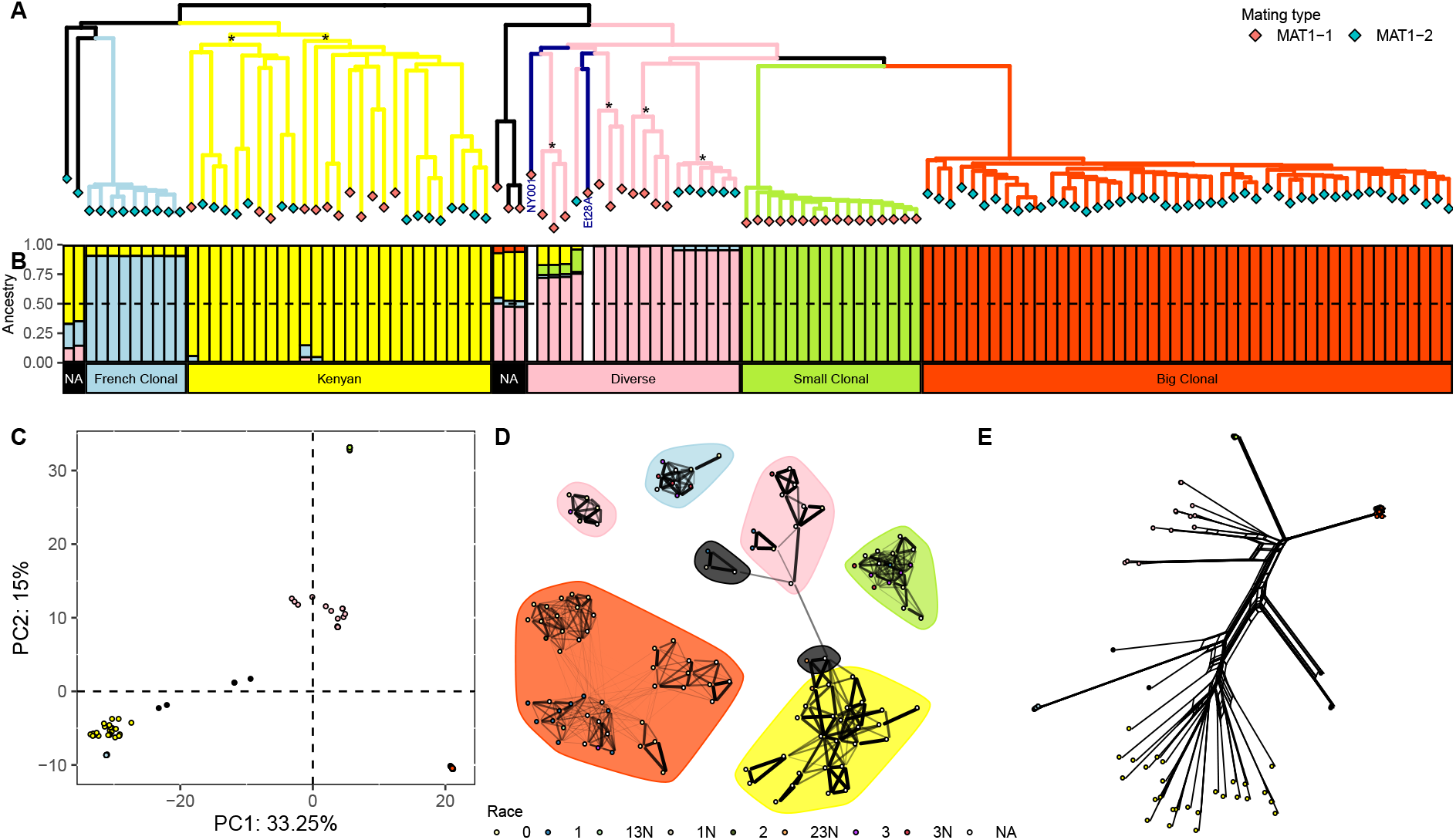
A) Rooted Neighbor-Joining tree from the polarized SNP dataset, branches are colored according to the observed ADMIXTURE clusters, dark blue indicates the two reference genomes (NY001 and Et28A), * sign indicates the subclusters within the major clusters (two subclusters in Kenyan and four in the Diverse cluster). Rhombuses in tip nodes are colored according to the mating type. B) Individual ancestry coefficients from ADMIXTURE for K=5 in the same order as the rooted NJ Tree. White gaps correspond to the two reference genomes which were not analysied in ADMIXTURE. NA show admixed individuals with no cluster assigned. C) First two axes of a PCA colored according to the five observed ADMIXTURE clusters. D) Population network created with CONE colored according the phenotyped race (NA in white for unknown race). Background color highlights the five ADMIXTURE classification clusters. E) Neighbor-Net created with SplitsTree colored according to the five observed ADMIXTURE clusters.

We also observed genetic differentiation within clusters. ADMIXTURE identified two distinct subclusters within the Kenyan cluster (*K* = 7), the Diverse cluster (*K* = 6), and the Big Clonal cluster (*K* = 8; Supplementary Fig.S1). CONE identified four connected subclusters within the Big Clonal cluster and two disconnected subclusters in the Diverse cluster (Fig.1D). The latter may consist of distinct clonal lineages that originated by recombination as shown by the Neighbor-Net. In contrast, no recombination is evident within the Big Clonal cluster and its subclusters, which therefore reflect evolutionary lineages of independent mutations.

To test whether the four European genetic clusters are geographically clustered, we analysed the spatial autocorrelation with Moran’s *I* using ADMIXTURE ancestry coefficients (*K* = 5; Fig.2A). Correlograms of Moran’s *I* indicate a wide geographic distribution and absence of geographic clustering of the Diverse and Big Clonal clusters (Fig.2B). In contrast, the French Clonal cluster is strongly clustered in France and the Small Clonal cluster at sampling locations within and between the Upper Rhine Valley and the border between Northwestern Austria and Southeastern Germany.

**Fig 2.**
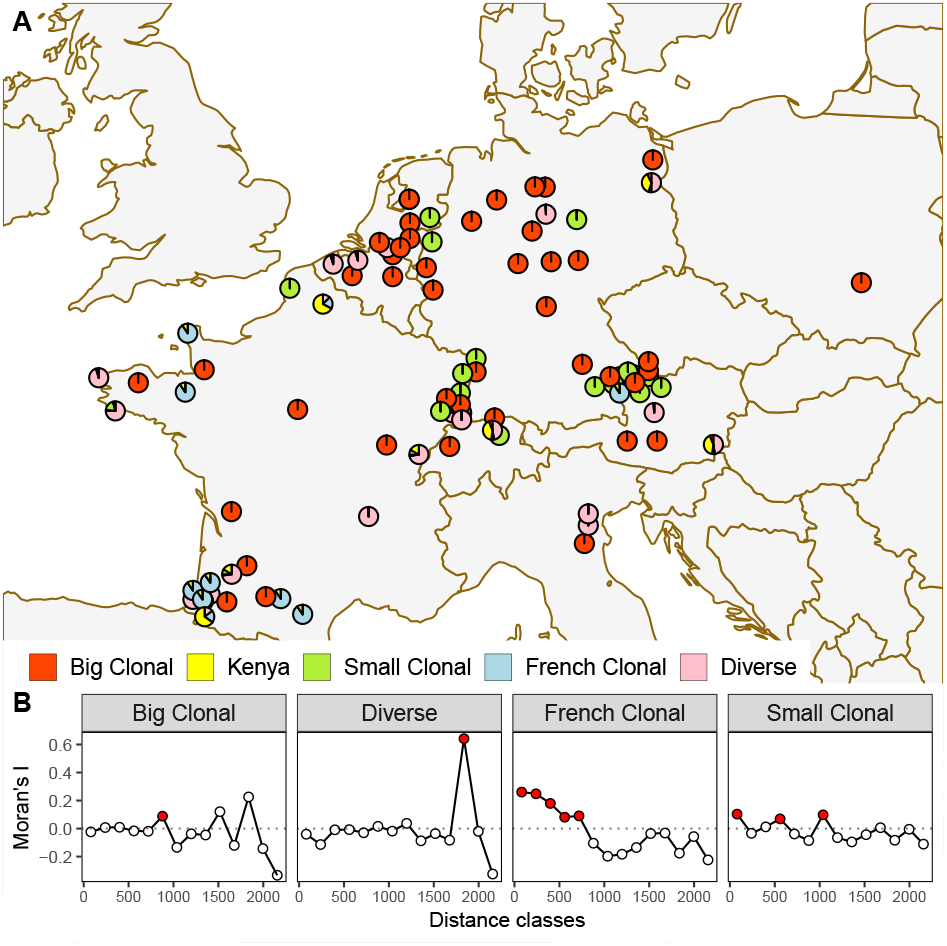
A) Geographic origin of European isolates. Pie charts indicate ancestry coefficients for *K* = 5 to show the geographic distribution of the five major genetic clusters. Geographically close isolates are shifted to avoid overlaping of pie charts. B) Correlogram of Moran’s *I* of the European ancestry coefficient along different distance classes. In red, *p*-value of Moran’s *I* < 0.05

### Mating type and recombination

Sexual reproduction in *S. turcica* is controlled by the *MAT1* locus with the *MAT1-1* and *MAT1-2* ideomorphs (Nelson, 1996;Turgeon, 1998), which are highly dissimilar alleles. Sequence reads from *MAT1-2* isolates do not map to a *MAT1-1* reference (and vice versa) resulting in an alignment gap. To determine the mating type of isolates we assembled all unmapped reads *de novo* into contigs and compared them with BLAST to a database of *S. turcica* sequences that included both *MAT1-1* and *MAT1-2* alleles. Isolates were classified as either *MAT1-1* or *MAT1-2* because all reads and contigs mapped to only one of the two mating types. Three clusters (Big Clonal, Small Clonal and French Clonal) are fixed for one mating type, whereas the Kenyan and Diverse clusters each have approximately 1:1 ratios of the two mating types, consistent with a history of sexual reproduction (Table 1). The presence of different mating types as indicator of sexual reproduction is supported by the Phi recombination test, which identifed past recombination events in the Kenyan and Diverse, but not in the other three clusters (Table 1). The test also detected recombination within the two Kenyan subclusters, of which each harbors both mating types in roughly equal proportions (Fig. 1A). We found no recombination within the four lineages of the Diverse cluster, consistent with the fixation of one mating type within each lineage of this cluster. A permutation test on the standardized index of association, 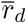, rejected the null hypothesis of random association of alleles in all five clusters, suggesting that despite past episodes of sexual reproduction, the Diverse and Kenyan clusters also show high rates of asexual reproduction in recent time.

**Table 1.** Diversity and reproduction type statistics of Kenyan and European isolates.

| Cluster | $n$ | $S$ | $\pi$ | $\theta_W$ | Tajima's $D$ | $MAT1-1 : MAT1-2$ | $p$-value | |
| --- | --- | --- | --- | --- | --- | --- | --- | --- |
| | | | | | | | Phi | $\bar{r}_d$ |
| Kenya | 26 | 11,880 | $6.631 \times 10^{-5}$ | $7.852 \times 10^{-5}$ | -0.62 | 11 : 15 | 0.0000 | 0.001 |
| Europe | 94 | 14,094 | $5.365 \times 10^{-5}$ | $6.949 \times 10^{-5}$ | -0.78 | 29 : 65 | - | - |
| Small Clonal | 16 | 393 | $1.52 \times 10^{-6}$ | $2.99 \times 10^{-6}$ | -2.15 | 16 : 0 | 0.4873 | 0.001 |
| French Clonal | 9 | 215 | $1.47 \times 10^{-6}$ | $2 \times 10^{-6}$ | -1.38 | 0 : 9 | 0.1034 | 0.001 |
| Diverse | 17 | 5,631 | $4.445 \times 10^{-5}$ | $4.201 \times 10^{-5}$ | 0.25 | 10 : 7 | 0.0000 | 0.001 |
| Big Clonal | 47 | 1,514 | $3.11 \times 10^{-6}$ | $8.65 \times 10^{-6}$ | -2.36 | 0 : 47 | 0.9229 | 0.001 |

Diversity statistics: $S$ : number of segregating sites, $\pi$ : nucleotide diversity per bp, $\theta_W$ : Watterson's estimator per bp, $D$ : Tajima's $D$ . Results are rounded to the number of presented digits. Reproduction type statistics: $MAT1-1$ : $MAT1-2$ as matying type counts. Phi ( $p$ -value): $p$ -value of the Phi recombination test. $\bar{r}_d$ ( $p$ -value): $p$ -value of the standardized test of random association of alleles.

The geographic distribution of the two mating types is correlated with the geographic distribution of the four European clusters as the mating type is fixed within each of the three European clonal lineages. However, mating types of the Diverse cluster are unequally distributed with a higher proportion of *MAT1-1* in the Southeastern part and a higher proportion of *MAT1-2* in the Northwestern part of its sampling area (Supplementary Fig.S2)

### Differences in genetic diversity between clusters

Consistent with their different histories of sexual and asexual reproduction, the five clusters also differ by their level of nucleotide variation (Table 1, Fig. 3A). Nucleotide diversity, π and Watterson’s estimator, *θ_W_*, are higher among the 26 isolates from Kenya (Genome-wide π = 6.631 × 10^-5^, per base pair) than among the 94 isolates from Europe (5.365 × 10^-5^). Both clusters harbor a high proportion of SNPs not present in the other cluster because only 4,647 SNPs segregate in both clusters, corresponding to 33% and 39% of the SNPs of the Kenyan and European clusters, respectively.

**Fig 3.**
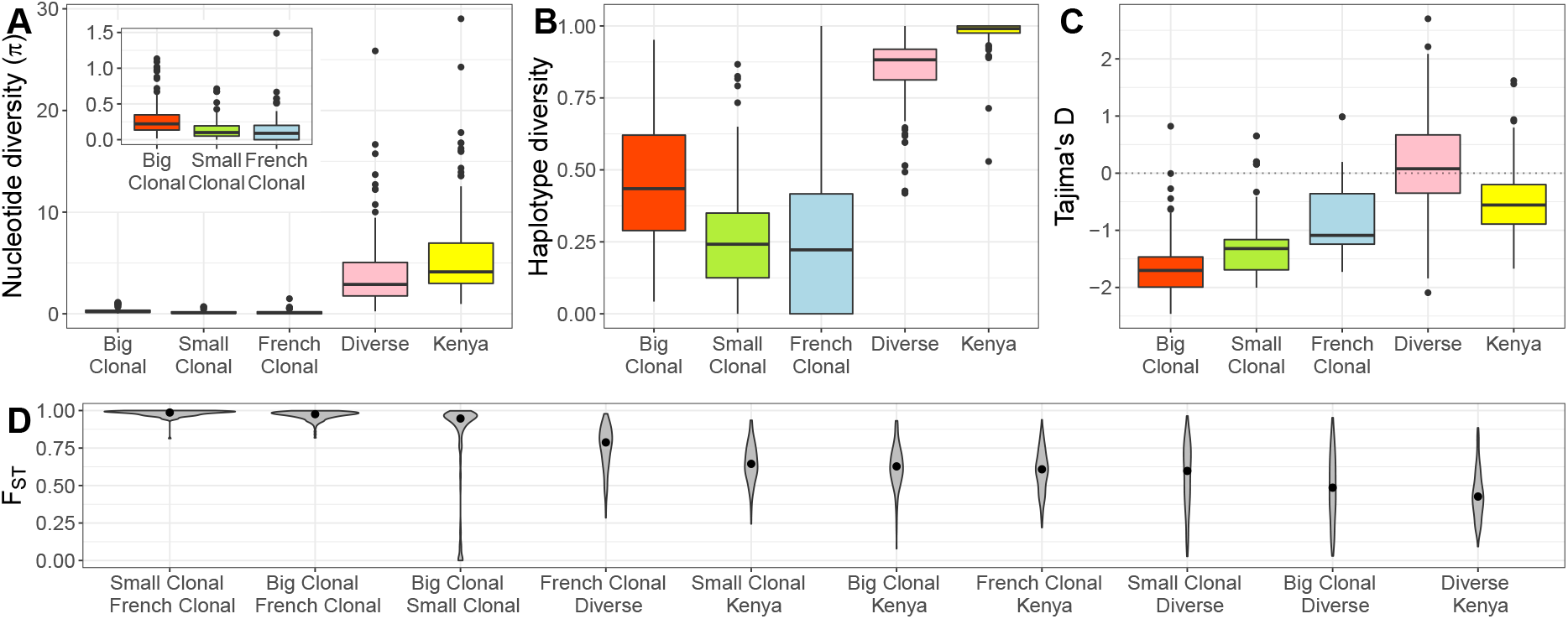
Levels of genetic diversity in five different genetic clusters defined by ADMIXTURE *k* = 5. A) Nucleotide diversity π per bp (in units of 10^-6^), B) haplotype diversity, C) Tajima’s *D*, and D) pairwise *F_ST_* calculated in windows of 250kb. The inset plot in (A) zooms into the *y* axis for the three first clusters (same units).

SNP-based genetic diversity differs between the four European clusters (Table 1A). The Diverse cluster shows 10 to 30 fold higher genetic diversity compared to the three clonal lineages. Its genetic diversity is 82% of the total European and 65% of Kenyan samples, respectively. Similar differences between the clusters are observed with haplotype diversity (Fig.3B). Tajima’s *D* values of the Big and Small Clonal clusters are highly negative (< 2) and less negative in the French Clonal cluster (−1.4; Table 1, Fig. 3C). The negative Tajima’s *D* values of the clonal lineages indicate a genome-wide excess of rare alleles that may be caused by demographic effects like population growth following recent emergence or genome-wide purifying selection. The first explanation was proposed for similar patterns in clonal lineages of other plant pathogens (e.g.,Latorre et al., 2020). Genetic differentiation of SNPs was measured as *F_ST_* and was highest between clonal lineages and smaller between the clonal lineages and the Diverse and Kenyan clusters, respectively (Fig. 3D).

The five clusters also differ in the distribution of genetic diversity along the genome. The Big Clonal, Small Clonal and French Clonal clusters have numerous genomic regions devoid of any genetic variation, whereas variation is more uniformly distributed in the Diverse and Kenyan clusters (Supplementary Figs. S3, S4 and S5). The lack of diversity is particularly strong for the Small Clonal and French Clonal clusters, because only 3% (Small Clonal) and 1% (French Clonal) of all 100 kb windows on the 15 longest scaffolds of the reference genome segregate for five or more SNPs. For the clonal lineages, most windows reflect the genome-wide negative Tajima’s *D* values and there are no visible outliers with highly negative Tajima’s *D* values that may reflect strong localized selective sweeps (Supplementary Fig.S6).

In contrast, both the Big and Small Clonal clusters have windows with highly positive Tajima’s *D* values (e.g. on scaffolds 4 and 10), which may indicate mapping errors caused by structural variants or strong balancing selection (Supplementary Table S2). However, these regions contain only very few (≤ 5) SNPs and, for the Big Clonal cluster, outlier Tajima’s *D* values do not deviate significantly from a neutral model of a constant or exponentially growing population (Supplementary Text A, Table S3).

### Tests of selection

To investigate whether the genetic clusters were affected by positive or purifying selection, we applied the McDonald-Kreitman (MK) test and compared synonymous and non-synonymous variation among isolates relative to the reference genome Et28A (Table 2). Although ratios of non-synonymous and synonymous substitutions (*D_n_/D_s_*) are frequently used for interspecific comparisons, they can also be interpreted for well separated clonal lineages (Kryazhimskiy and Plotkin, 2008). The Et28A reference clusters with the Diverse cluster, thus they are not well separated and we did not perform the analysis for the Diverse cluster. The ratio of synonymous to non-synonymous nucleotide diversity (*π_N_/π_S_*) estimates the fraction of effectively neutral mutations among all mutations (Akashi et al., 2012) under Ohtas’s nearly neutral model (Ohta, 1973). The Big Clonal, French Clonal and Kenyan clusters show *π_N_/π_S_* ratios below 1 indicating that a majority of mutations are non-neutral or nearly neutral. Variation in the Small Clonal cluster differs from a nearly neutral model with a ratio *π_N_/π_S_* = 2.1 and a much higher ratio of non-synonymous to synonymous mutations, *P_n_/P_s_* = 5.5 than the other clusters (Table 2). However, with the exception of the Small Clonal Cluster (p < 0.0001), a MK test does not reject the null hypothesis of neutral evolution indicating that purifying selection has no significant effect on the fate of mutations in four of the five genetic clusters of our sample, which is unexpected given the *π_N_/π_S_* ratios observed.

**Table 2.** McDonald-Kreitman test

| Cluster | $\pi_N/\pi_S$ | $P_n/P_s$ | $D_n/D_s$ | $NI$ | $p$-value |
| --- | --- | --- | --- | --- | --- |
| Big Clonal | 0.747 ($1.30 \times 10^{-6}/1.80 \times 10^{-6}$) | 1.648 (201/122) | 1.366 (168/123) | 1.206 | 0.283429 |
| Small Clonal | 2.123 ($8.00 \times 10^{-7}/4.00 \times 10^{-7}$) | 5.455 (60/11) | 1.480 (182/123) | 3.686 | 0.000055 |
| French Clonal | 0.299 ($4.00 \times 10^{-7}/1.30 \times 10^{-6}$) | 2.818 (31/11) | 1.653 (390/236) | 1.705 | 0.185753 |
| Kenya | 0.506 ($2.47 \times 10^{-5}/4.88 \times 10^{-5}$) | 1.734 (1278/737) | 1.395 (113/81) | 1.243 | 0.161458 |

$\pi_N/\pi_S$ is the ratio of non-synonymous to synonymous nucleotide diversity; $P$ : population polymorphisms; $D$ : fixed derived mutations (reference Et28A as outgroup); $n$ : non-synonymous mutations, $s$ : synonymous mutations; $NI$ : Neutrality Index, calculated as $(P_n/P_s)/(D_n/D_s)$ ; $p$ -value: Fisher's exact test $p$ -value.

### Inference of split times

To investigate the demographic history of European isolates we included the two North American isolates Et28A and NY001 and used the polarized SNP data.A rooted tree revealed a close relationship of the American and European isolates (Fig.1A), which was independently confirmed by merging our resequencing data with genotyping by sequencing (GBS) data of 13 North American isolates (Mideros et al., 2018) resulting in a set of 280 genome-wide SNPs (Supplementary Fig.S7). The resulting phylogenetic tree and PCA plot (Supplementary Fig.S7) of the merged dataset are essentially identical to the analyses of European isolates based on the complete sequencing data. Both methods group the North American isolates with the Diverse cluster, consistent with the tree in Fig.1A.

To test whether European clonal lineages split before or after their introduction to Europe we estimated divergence times between the five clusters as time back to the most recent common ancestor (MRCA) of a pair of clusters, and emergence times of clonal lineages within clusters as time back to the MRCA within in each cluster using BEAST (Fig.4A and Supplementary Fig.S8). The three clonal clusters diversified quite recently with posterior mean emergence of the most recent common ancestor in the year 1985 for the Big Clonal (1978-1990 include ≥ 95% posterior mass with highest posterior density; 95%HPI), 1998 for the Small Clonal (1993-2001; 95%HPI) and 1999 for the French Clonal (1995-2002; 95%HPI) clusters. Split times between clusters are more distant and range from the year 1609 between Small Clonal and Big Clonal (1480-1809; 95%HPI), 1503 between the ancestors of Big Clonal, Small Clonal and the Diverse cluster (1456-1667; 95%HPI) to 1198 between the ancestors of the Big Clonal, Small Clonal, North American reference isolates and French Clonal (975-1368; 95%HPI). The Diverse cluster emerged much later than the clonal clusters in 1520 (1386-1624; 95%HPI) and the nodes of its genealogical tree are more spread over time. Split times and tree topology of the BEAST analysis agree with the phylogeny in Fig.1A and support a much closer pairwise relationship of Big Clonal and Small Clonal than to the French Clonal cluster. Including non-clonal lineages in analyses to estimate split and emergence times may introduce a bias due to reticulate events (Latorre et al., 2020). For this study, there is no meaningful bias introduced (Supplementary Text B).

**Fig 4.**
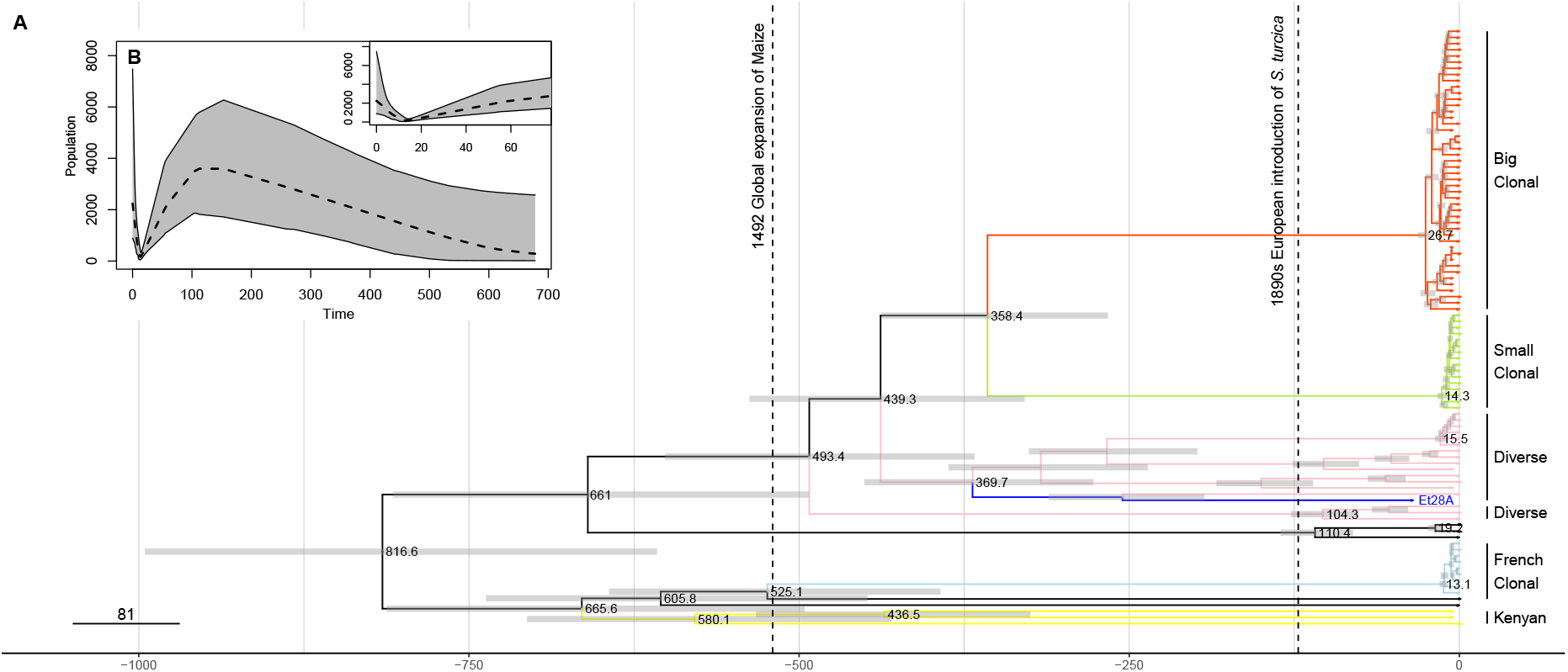
A) Dated phylogeny obtained with BEAST, using all European isolates, reference genome Et218A (in dark blue) and three samples from Kenyan cluster. Time is given as years before 2012, the year of the most recent sampling. Horizontal gray bars show the 95 % highest posterior density intervals (95% HPI) for split times. B) Extended Bayesian skyline plot obtained with BEAST for the analysis from (A). The inset zooms into the most recent past. Time runs backwards from 2012. The dashed line shows the median posterior population size, while the gray area shows the 95% HPI.

We then investigated whether the global expansion of maize cultivation after the beginning of the Columbian exchange in 1492 and the strong increase of maize cultivation in Europe during recent decades was accompanied by an increase of the effective pathogen population size, *N*_e_. After adding global population size as parameter to the phylogentic model for BEAST median, posterior estimates of *N*_e_ changed substantially over time (Fig. 4B and Supplementary Figure S8). Estimates of *N*_e_ based on the European samples, three samples from Kenyan cluster and the North American reference sequence indicate a long phase of population growth since the time of the most recent common ancestor (MRCA) of the European samples about 825 years ago until a period between 1859 and 1900, followed by a population decline until 1999, when population size *N*_e_ was lower than at the time of the MRCA. This decline was then followed by a very recent epoch of strong population growth for 20 years until the last sampling date 2012. A recent, rapid growth is consistent with strongly negative genome-wide Tajima’s *D* values within the three European clonal clusters. A decline of *N*_e_ followed by recent strong growth was confirmed by analysing only Big Clonal, Small Clonal, and French Clonal clusters together with the reference genome (Supplementary Figs. S9 and S10).

### Neutral versus selection-driven population dynamics

The low genetic diversity and genome-wide excess of rare polymorphisms within clonal lineages may reflect rapid population growth or result from recurrent, short phases in which newly emerged genotypes with a skewed offspring distribution become dominant. Among predominately asexually reproducing fungal pathogens, following processes may lead to a skewed offspring distribution even without population size changes: (*i*) rapid selection of newly emerged genotypes with a very high fitness coefficient (Neher and Hallatschek, 2013;Latorre et al., 2020), (*ii*) a large number of offspring originating by chance from a single parental genotype analogous to sweepstake reproduction in marine species (Steinrücken et al., 2013;Dutta et al., 2020), or (*iii*) a large number of offspring from genotypes that evolved virulence against monogenic resistance genes present in maize varieties (boom-bust cycles) (Tellier and Lemaire, 2014). Genealogies in these cases can be modeled as multiple-merger coalescents. We compared these models with a standard Wright-Fisher type reproduction with growing population sizes, modeled via a bifurcating Kingman coalescent with exponential growth. For the Big Clonal and Kenyan clusters we compared both coalescent models using SNPs segregating within this cluster on the five largest scaffolds of the current *S. turcica* reference genome using a Random Forest Approximate Bayesian Computation (RF-ABC) approach for model comparison and parameter estimation. In the other clusters we observed high prior error rates and low posterior probabilities and considered these results not as robust (Supplementary Table S5). Table 3 shows the results of the RF-ABC analysis. For the Big Clonal cluster, it provides strong support for a bifurcating Kingman coalescent with strong exponential growth over a multiple merger coalescent, which refutes strong selection without growth or sweepstake reproduction. The Kingman coalescent is also preferred for the Kenyan cluster, but with much smaller growth rates. All scaffolds in the Big Clonal and in the Kenyan cluster show ‘positive’ (≥ 3 odds ratio) to ‘strong’ support (≥ 20 odds ratio, only for Big Clonal) for an exponential growth model over a multiple merger model according to the Kass-Raftery scale (Kass and Raftery, 1995). Simulations showed that the observed genetic diversity in both the Kenyan and Big Clonal clusters are obtained with the best-fitting model (Supplementary Text C, Figs. S11 and S12). Although our analyses reject the hypothesis that skewed offspring distribution alone shapes genetic diversity for the Big Clonal and Kenyan clusters, a combination of exponential growth with skewed offspring distributions explains the data similarly well as neutral population growth (Supplementary Text D).

**Table 3.** RF-ABC model selection results

| Cluster | SNP | Mean | Post. prob. | Odds | Fitted |
| --- | --- | --- | --- | --- | --- |
| | count | OOB | Exp. growth | ratio | rate $g$ |
| Big Clonal | 92-128 | 21-22% | 83 - 98% | 4.9-49 | 181-630 |
| Kenya | 683-1007 | 20% | 75.2 - 84.3% | 3-5.4 | 5.5-8.5 |

Mean OOB is the out-of-bag prior error rate for model classes, averaged over all model classes. Post. prob. exp. growth gives the posterior probability of the exponential growth model. Fitted parameter $g$ : Posterior median of the exponential growth parameter in coalescent units, where one unit represents $2N$ generations. For each variable, we report the range of values for the five biggest scaffolds. For the transformation of posterior probabilities into odds ratios for a comparison of exponential growth vs. any other genealogy model used, see Materials and Methods.

### Different pathogen races within clonal lineages

To test whether isolates within clonal lineages belong to the same or different races, we identified 62 isolates in our sample whose race was determined in a race monitoring of 542 European *S. turcica* isolates collected in 2011 and 2012 from the major maize growing regions in Europe (Hanekamp, 2016). The monitoring revealed that race 0 was the most dominant with 45% of isolates, followed by race 1 (22%), 3 (15%) and 3N (14%). Only 4% of isolates were virulent against two or more resistance genes (races 13, 123, 23, 2, 23N, 12, 1N and 13N). Mapping the race type of the 62 isolates onto the CONE network reveals that Big and Small Clonal clusters harbor four races each and the French Clonal cluster three races (Fig. 1D). Single, independent *de novo* mutations in pathogen effector genes are sufficient to create new races and may explain the diversity of races within clonal lineages. Alternatively, the presence of the same races in different lineages may reflect shared polymorphisms that originated in ancestral populations although such an explanation seems unlikely given the low genetic diversity within clonal lineages (Table 1).

### Identification of divergent regions and structural variants

The differences in SNP allele frequencies between clonal lineages suggests that highly divergent genomic regions and presence absence structural variants (PAVs) also contribute to genomic differentiation. We therefore used sequence read coverage and *k*-mer frequencies to identify highly divergent regions and PAVs. First, we calculated for each isolate its sequence coverage of the reference genome in 43,443 windows of 1 kb length and expressed coverage as percent bases covered by at least one sequence read in each window. Windows with low coverage indicate a high proportion of mapping gaps in the reference and windows with a highly variable coverage between isolates pinpoint structural variants. Using the top 2.5% windows (*n* = 1, 012) with the most variable sequence coverage between isolates, we constructed a NJ tree from a pairwise Euclidean distance matrix of reference sequence coverage (Supplementary Fig. S13) to cluster isolates with similar variation in coverage. The topology of the resulting tree is highly similar to the rooted SNP-based tree indicating that highly variable regions and PAVs reflect similar genealogical process than SNP allelic variation.

To identify genomic regions that differentiate pairs of clonal clusters we used HAWK (Rahman et al., 2018), which identifies *k*-mers whose frequency differs between clusters. Among all pairwise comparisons (see Methods), we obtained different *k*-mer frequencies only between the Big Clonal vs. Small Clonal clusters. Among 6,341 *k*-mers that differentiate the two clusters, 3,048 are associated with the Big and 3,293 with the Small Clonal cluster. We de novo assembled both *k*-mer clusters independently into longer sequence contigs and found that 93% of assembled *k*-mers mapped to few, distinct regions of the reference genome, suggesting that a small number of genomic regions contribute to genomic differences between the two clusters. Assembled *k*-mers mapped to only 1,167 (12.73 %) 10kb windows of the reference genome. There were only 30 windows (0.33% of all windows) that collected the top 2.5% *k*-mer counts with at least 9.85 mapped *k*-mers per window. Among all mapped *k*-mers, 25% map to these 30 windows, which tend to be highly repetitive. A majority of 22 out of 30 windows (73%) is highly repetitive with ≥ 50% repetitive elements and no window contains gene-rich regions.

To identify proteins that may differentiate the Big and Small Clonal clusters, we conducted a BLASTX analysis against a non-redundant BLAST protein database with the remaining unmapped *k*-mers. For both clusters, ‘hypothetical protein’ was the most frequent annotation of proteins among the five best hits with a cutoff e-value of < 0.001, followed by the mating type *MAT1-2* for the Big Clonal cluster. The latter finding is a positive control of the *k*-mer mapping approach because Big Clonal has *MAT1-2*, which does not map to the reference genome Et28A, and Small Clonal has *MAT1-1*, which maps to the reference genome. For the Small clonal cluster, the second most frequent BLAST hit was ‘polyketide syntase protein’, which is potentially associated with pathogen virulence (Ohm et al., 2012). We also used the race assignment to identify *k*-mers associated with race-specific genes, however no significant and robust outcome was found (Supplementary Table S6). This negative result may either reflect a too small sample size or genetic differences of single or few variants that are not uncovered by the analysis of *k*-mers.

## Discussion

Our work confirms earlier studies of *S. turcica* genetic diversity and mode of reproduction in Europe and Africa (Borchardt et al., 1998,?). Isolates originating from Kenya form a single cluster with high genetic diversity, equal frequency of both mating types and genomic patterns of recombination consistent with a higher rate of sexual reproduction of *S. turcica* in tropical climates. In contrast, European isolates are composed of four distinct clusters, which differ by their relative frequency and geographic distribution. Three clusters (Big Clonal, Small Clonal, French Clonal) represent single clonal lineages, whose genetic diversity is very low, do not show evidence of recent recombination, and are fixed for one of the two mating types. The fourth cluster (Diverse) consists of diverse clonal lineages that, taken together, have a high level of genetic diversity, evidence of past recombination and an equal frequency of both mating types. These three characteristics in combination with low *F_ST_* values between the Diverse vs. the Big Clonal and Small Clonal clusters, respectively, suggest that the Diverse cluster is a source of genetic diversity from which clonal lineages emerged previous to the arrival of *S. turcica* in Europe. The two North American isolates Et28A and NY001 cluster with the Diverse cluster and are highly similar to different European isolates indicating the close connection between European and American samples that may reflect an American origin of the Big Clonal, Small Clonal and Diverse clusters. In contrast, the French Clonal cluster is closely related to the Kenyan cluster and therefore likely has the same origin. A previous study interpreted the presence of African alleles in an isolate from Southwestern France of *S. turcica* as recent migration (Borchardt et al., 1998). This is not supported by our analysis because the split time of Kenyan and French Clonal clusters predates the arrival of *S. turcica* in Europe, and their close relationship reflects a common ancestry instead of recent migration.

Our divergence time estimates suggest that individual lineages within each of the three Big Clonal, Small Clonal and French Clonal clusters emerged less than 40 years ago. Although these very recent emerge times are based on a limited sample we consider them reliable (Saunders et al., 1984) (Supplementary TextE). In contrast, the divergence times of the five clusters identified in our sample are more distant and range between 816 to 360 years ago (Fig.4), which predates the introduction of *S. turcica* into Europe and strongly suggests these clusters originated outside Europe and were independently introduced. The clonal sublineages within the Diverse cluster originated between 370 to 50 years ago and are separated by sexual recombination events, which are unlikely under European climatic conditions. For this reason, they were likely independently introduced into Europe.

### Evolutionary forces determining pathogen demography

Although multiple crop pathogens expanded globally in short time, only few studies analysed the evolutionary forces determining expansions, in particular the role of selection on plant pathogens, using explicit population genetic modeling (Croll and McDonald, 2017). We employed Approximate Bayesian Computation (ABC) to compare two coalescent models and to differentiate between a neutral model of exponential growth and a selection-based model of the *S. turcica* expansion in Europe. Simulations of models with asexual reproduction demonstrate a high power of ABC to differentiate between neutral and selection-driven demographies with suitable summary statistics (Freund and Siri-Jé gousse, 2020). The Big Clonal cluster is particularly interesting for such an analysis because it is the most successful cluster in terms of sample frequency and geographic distribution in Europe. Its large sample size provides better statistical power and a restriction of ABC to clusters without a strong internal population structure removes a bias in distinguishing among genealogy models (Koskela and Wilke Berenguer,2019). The ABC analysis of the Big Clonal cluster (Table 3) reveals a recent population size increase and in addition that observed genetic diversity in this cluster is not consistent with a history of rapid selection as proposed for *Magnaporthe oryzae* (Latorre et al., 2020) or boom-bust cycles caused by host-pathogen coevolution without population growth. In other fungal crop pathogens such as *Zymoseptoria tritici* random fluctuations in fecundity and a potential for very large offspring numbers per individual have been proposed (Dutta et al., 2020), which should lead to a multiple merger genealogy if it is strong enough. Our results also exclude such a model as sole explanation for the observed diversity in the Big Clonal Cluster or indicate that fecundity differences in the pathogen are too small to affect the shape of the genealogy. Instead, observed diversity within this cluster can be explained by just assuming neutral population growth. These analyses do not exclude the possibility that an exponential increase of *N_e_* in the Big Clonal cluster results from a relative fitness advantage caused by adaptive *de novo* mutations or a favourable combination of adaptive mutations achieved via sexual recombination in the founders of the cluster. In addition, a more complex pattern of neutral population growth on top of selection processes or sweepstake reproduction can also not be ruled out. Since the Kenyan cluster also supports a neutral coalescent model with a low rate of population growth there is no reason to expect multiple mergers as standard gene genealogies in *S. turcica*. The absence of interpretable results for the ABC analyses for the Small Clonal and French Clonal clusters likely results from too small sample sizes.

Tests of neutrality based on comparisons of non-synonymous and synonymous genetic diversity (Table 2) do not contradict a model of neutral evolution as main driver of genetic diversity for the Big Clonal, French Clonal and Kenyan clusters. Although these clusters exhibit an excess of non-neutral diversity, there is no strong selection against it as indicated by non-significant MK test results. This seems contradictory, but may result from (*i*) clonal interference, which prevents efficient selection against deleterious mutants, or (*ii*) from a surplus of beneficial founder mutations that offsets the effect of purifying selection, although this requires many beneficial mutations and is therefore unlikely, or (*iii*) reflect an underpowered MK test. Nevertheless, this result shows that purifying selection is not a major determinant of genetic diversity within these clusters. The Small Clonal cluster may have a different evolutionary trajectory because the significant MK test result for purifying selection contrasts an excess of non-synonymous diversity, which suggests that evolution in this cluster does not follow a nearly-neutral model. Overall, *S. turcica* genetic clusters do not provide evidence for strong purifying selection, which is in contrast to the rice fungal pathogen *Magnaporthe oryzae* (Gladieux et al., 2018). Lack of evidence for selection as main determinant of genetic diversity in the Big Clonal cluster and a temporal coincidence of *S. turcica* population growth with the expansion of maize cultivation in Europe leads us to propose that the demographic expansion of this lineage was not driven by rapid evolutionary adaptation to European maize varieties or the environment.

### Limited evidence for host-pathogen co-evolution in Europe

Our sample of isolates was collected in 2011 and 2012 and represents a snapshot in time and space that is restricted to Europe and Kenya. Both factors limit further interpretations of our results and lead to questions about the role of *S. turcica* - maize coevolution within and outside Europe. First, the demographic analysis suggests an independent and recent single introduction of the French Clonal, Small Clonal and Big Clonal clusters into Europe (Table 3), although our results are also consistent with independent, repeated introductions of the same clonal lineages. For example, the clonal lineages within the Diverse cluster originated by sexual recombination over an extended period of time (Fig. 4). Since sexual recombination is unlikely under European climatic conditions, the lineages likely originated outside Europe and were then subsequently introduced. Further evidence for repeated introductions is the high genetic similarity among European and North American isolates suggesting recent exchange or a common origin in a different region, such as Mexico, because European isolates were more similar to Mexican than to Kenyan isolates (Borchardt et al., 1998). Additional samples from putative regions of origin such as Central America and tropical Africa are required to resolve this issue.

A second question refers to the effects of maize resistance genes on *S. turcica* evolution and epidemiology. There is no association between the five genetic clusters and the distribution of *S. turcica* races among these clusters. This observation and a high proportion of race 0 (i.e., non-virulent against four tested *Ht* genes) isolates in all five clusters shows that race-specific virulence did not generate new pathogen lineages with a strongly increased fitness. In combination with the evidence for neutral evolution of genetic variation in the European isolates, we conclude that strong selection against qualitative or quantitative maize resistances had very little or no effect on genetic diversity in Europe. However, future studies should associate the genetic diversity of host and pathogen genomes using joint association analysis, (e.g.,Wang et al., 2018), to elucidate the role of genotype by genotype (GxG) effects in the spatial and temporal dynamics of host-pathogen interactions. Such information will contribute to avoid breakdown of resistance genes and achieve long-term resistance management (Nelson et al., 2018).

A third question refers to the evolution or new races, because the presence of multiple races within the five *S. turcica* clusters suggests a rapid and repeated breakdown of *Ht*-based monogenic resistances in maize varieties (Fig.1D). Since selection against host resistance does not seem to affect the evolutionary dynamics of *S. turcica*, a frequent origin of new races may be facilitated by a high mutation rate, which we estimated as posterior mean substitution rate of 10^-4^substitutions per year per site using BEAST. This rate is much higher than in *Magnaporthe oryzae*, where it was estimated to be in the order of 10^-8^ (Gladieux et al., 2018). A *k*-mer based association analysis did not find *k*-mers that are significantly associated with race type, possibly because of small sample sizes, whereas *k*-mer analysis of the complete sample identified the mating type gene and several genomic regions that differentiate clonal groups and harbor genes with putative roles in pathogenicity.

In conclusion, our analyses indicates a rapid spread of different *S. turcica* clonal lineages in Central and Western Europe in the absence of both recombination and strong selection for pathogen virulence. Monitoring of pathogen diversity on larger geographical scales and over time is required to fully understand forces influencing pathogen epidemiology and evolution, and the evolution of pathogen races. However, our work shows that large scale sequencing and population genomic analysis provide useful information to develop breeding programs informed by host-pathogen evolution and to control plant pathogens by improved agricultural management.

## Materials and Methods

### Cultivation of fungal isolates

The origin and sampling information of isolates is described in Supplementary Dataset S1. Lyophilized isolates were transferred to Becton Dickinson BBD Potato Dextrose Agar plates and incubated for at least 10 days at 25°C and a 12h light / 12hr dark cycle until plates were completely covered by mycelia. This fungal tissue was scraped from the surface with a spatulum and collected in a 2 ml plastic reaction tube.

### DNA extraction and NGS sequencing

After adding six ceramic beads (2.8 mm diameter; MoBio, USA) to each tube, the tissue was ground in a Retsch mixer mill (MM400) for 30 sec at a speed of 30 sec^-1^. The DNA was then extracted with the Micro AX Blood Gravity KI (A&A Biotechnology, Poland; Cat. No. 101-100) according to manufaturer’s instructions and diluted to a concentration of 2.5 ng μl^-1^ EB buffer. Whole genome sequencing libraries were generated using a multiplex tagmentation protocol (Baym et al., 2015) with minor modifications. The assignment of barcodes to isolates is detailed Supplementary Dataset S1. The libraries were paired-end sequenced (2 x 100 bp) on a HiSeq 2500 Illumina sequencer (Macrogen, Korea) in three batches of 24, 96 and 46 isolates, respectively.

### Read mapping and variant calling

Raw Illumina reads processed with Trimmomatic v0.36 (Bolger et al., 2014) were mapped with BWA v0.7.12-r1039 (Li and Durbin, 2009) against the *Setosphaeria turcica* reference genome Et28A v1.0 (race 23N strain 28A) (Ohm et al., 2012;Condon et al., 2013) obtained from EnsemblFungi version 39 (Kersey et al., 2016). After removing PCR duplicates with Picard tools (http://broadinstitute.github.io/picard/) and realigned reads with GATK v3.7-0 (McKenna et al., 2010) samples that had low percentage of mapping reads (<83%) and/or low coverage (<5X) were excluded. For variant calling we kept bi-allelic SNPs discovered in both GATK and samtools/bcftools callers (Li et al., 2009; Li, 2011;Narasimhan et al., 2016). SNPs were filtered for minimum read depth of 3 and a maximum of 100, minimum proportion of reads supporting a genotype call of 0.8, maximum percentage of missing data per SNP of 35%. Missing genotypes were imputed with multiple correspondance analysis (MCA) using the R package `missMDA` (Josse and Husson, 2016).

### Variant polarization

To polarize alleles we used the reference genomes of two closely related species, *Bipolaris sorokiniana* ND90Pr (Ohm et al., 2012;Condon et al., 2013) and *Bipolaris maydis* ATCC 48331 (Ohm et al., 2012;Condon et al., 2013) that were both obtained from EnsemblFungi. Outgroup genomes were aligned with *Setosphaeria turcica* reference genome using TBA (Blanchette et al., 2004). Genotype calling from the alignment was done with MafFilter (Dutheil et al., 2014) and only bi-allelic variants shared with both outgroup species were kept. Ancestrality of the alleles was assigned to the allele of the outgroup species and genotypes of the 130 samples (129 isolates and *S. turcica* reference genome) were polarized accordingly. Additionally, we included the draft genome of *S. turcica* (race 1 strain NY001; JGI Fungal Program (Grigoriev et al., 2012;Nordberg et al., 2014) under GOLD Project ID Gp0110874), originally collected in Freeville, New York in 1983 (Chung et al., 2010) as additional sample to the polarized SNP dataset.

### Mating type assignment

The *S. turcica* Et28A reference genome is of mating type *MAT1-1*. For this reason, isolates with a mapping gap on the *MAT1* locus were candidates for *MAT1-2* type. Confirmation of the mating type was done with a *de-novo* alignment of the unmapped reads with MegaHit (Li et al., 2015) and posterior blasting with BLAST (Altschul et al., 1990;Camacho et al., 2009) to a nucleotide database of *S. turcica* (which included the sequences of MAT1-1 and MAT1-2).

### Population structure

To infer population structure we conducted Principal Component Analysis (PCA), maximum likelihood estimation of individual ancestries with ADMIXTURE (Alexander et al., 2009) and population network estimation with community detection using neighborhood selection with CONE (Kuismin et al., 2017). PCA was calculated with the glPca function of the `adegenet` (Jombart, 2008;Jombart and Ahmed,2011) R package. We used *K* = 5 clusters because higher values did not significantly reduce cross-validation error (Supplementary Fig.S14) and produced the same composition of clusters with *K* = 5 in 20 independent runs. The different admixture runs were merged with CLUMPAK (Kopelman et al., 2015). Population networks were estimated using R scripts for haploid data provided by CONE authors (Kuismin et al., 2017). An unrooted Neighbor-joining tree was built from Euclidean distances and Neighbor-net was calculated with SplitsTree v4.14.6 (Huson and Bryant, 2006) using Hamming distances calculated from the unpolarized SNP dataset. A correlogram on Moran’s *I* (Cliff et al., 1981) was used to test the European spatial autocorrelation along different distance classes of equal frequency using `correlog` function from the R package `pgirmess` (Bivand and Wong, 2018;Bivand et al., 2013). As quantitative variable we used the individual’s ancestry coefficient of ADMIXTURE *K* = 5.

### Diversity statistics

Numbers of segregating sites *S*, genome-wide nucleotide diversity π, Watterson’s estimator *θ_W_* and Tajima’s *D* were calculated for the two sets of European and of Kenyan isolates, as well as for the subpopulations within them. The single isolate from Turkey within the Kenyan cluster (WGRS-Test 23) has a strong effect on its diversity measures because it contributes 886 additional SNPs (7% of the total). Since this sample is geographically separated from Kenya, we excluded it from the subsequent analysis of the Kenyan cluster. Both π and *θ_W_* are reported per base pair by dividing genomewide values by the maximum number of bases aligned to the reference across all sampled isolates (which are 39,649,104 of 43,013,545). For the 15 biggest scaffolds, an additional sliding window analysis for windows of size 100k bp was performed. We computed π, *θ_W_*, Tajima’s *D* and the haplotype diversity per window for all groups with the R package `PopGenome` (Pfeifer et al., 2014).

### MK test

To calculate the McDonald–Kreitman (MK) test we first ran SnpEff version 4.3t (Cingolani et al.,2012) with the–*classic* output style and Setosphaeria turcica et28a genome version on vcf subsets that included i) only population polymorphisms and ii) only fixed derived mutations. We used reference Et28A as outcluster, and excluded positions were Et28A and the sample WGRS 62 (closest sample to Et28A) were different. Neutrality Index (*NI*) was calculated as 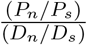, where *P* are polymorphisms, *D* substitutions, *s* synonymous mutations and *n* non-synonymous mutations. Fisher’s exact test *P*-value was computed using the 2×2 contingency table of the four type of mutations. π_N_/π_S_ ratio was calulated as

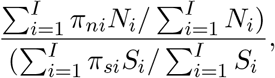

where *I* is the number of scaffolds, *N_i_* is the number of non-synonymous sites in scaffold *i*, *S_i_* the number synonymous sites in scaffold *i* and *π_ni_*, *π_si_* are the non-synonymous or synonymous nucleotide diversities per non-synonymous or synonymous site in scaffold *i*. All *π_ni_*, *π_si_*, *N_i_* and *S_i_* were obtained from *population_summary.txt* output file after running SnpGenie (Nelson et al., 2015).

### Analysis of reproduction type

The reproduction type (clonal vs. sexual) was analyzed with three approaches. First, we calculated the mating type ratio for each population. A 1:1 ratio of the mating type is a strong indicator for sexual reproduction whereas a significant skewed ratio indicates clonal reproduction (Sommerhalder et al.,2006;Milgroom, 1996). Second, we tested for recombination using the Phi test (Bruen et al., 2006) as implemented in SplitsTree and third, we tested the null hypothesis of random association of alleles by 999 permutation tests of the standarized index of association 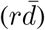 with `poppr` (Kamvar et al., 2014).

### Analysis of demography within clonal subpopulations

For the five clusters Big Clonal, Small Clonal, French Clonal, Diverse and Kenyan, we performed model selection between sweepstake reproduction using genealogies modelled by Dirac- and Beta-*n*-coalescents, or rapid selection using the Bolthausen-Sznitman *n*-coalescent in a fixed-size population, and standard reproduction with a Kingman’s *n*-coalescent in a fixed-size or an exponentially growing population. Additionally, we performed parameter estimation within the best-fitting model class. Model selection and parameter estimation was performed via random-forest based Approximate Bayesian Computation (Pudlo et al., 2015;Raynal et al., 2019) using quantiles of summary statistics for unpolarized data (Freundand Siri-Jé gousse, 2020). For the analysis, we treated each scaffold as a single non-recombining locus and ran it on the 5 biggest scaffolds. We considered Beta(2 - α, α)-*n*-coalescents with α ∈ (1, 2) where α = 1 denotes the Bolthausen-Sznitman *n*-coalescent, Dirac *n*-coalescents with parameter p ∈ (0, 1) and, for Kingman’s *n*-coalescent, exponential growth rates in (0, 2500). We set a uniform prior on *p* for Dirac-*n*-coalescents, while for Beta-*n*-coalescents, we set α = 1 with a probability of 5% and in all other case draw α uniformly from (1, 2). For Kingman’s *n*-coalescent with exponential growth, we used an uniform prior on the log scale on (0, 2500) with a spike corresponding to a prior probability of 0.02 at *g* = 0.

The scaled mutation rate *θ* was set to the generalized Watterson estimator *θ_W_* = 2*S/E*(*L_n_*), where *L_n_* is the expected total length of the underlying genealogy model, but with a random fluctuation around this estimate as in Freund and Siri-Jégousse (2020, Scenario 3). As statistics, we used the (.1,.3,.5,.7,.9)-quantiles of the branch length of the neighbor-joining tree reconstructed from the genetic data, of the Hamming distances and of the linkage disequilibrium statistic *r*^2^, as well as the number of segregating sites *S*, nucleotide diversity π and the folded site frequency spectrum, where all minor allele counts above 15 are summed up as a single statistic. Each model class is simulated 175,000 times and the random forest is built from 500 trees. Simulations were performed as in Freund and Siri-Jé gousse (2020), ABC parameter estimation and model selection were conducted with the R package `abcrf`. An estimated lower bound for odds ratio/Bayes factors of the best fitting model to any other model was given by 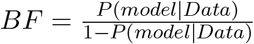, i.e. we treat the posterior probability of any other model as posterior probability that the best fitting model is not the true model.

### Phylogenetic dating with BEAST

We ran BEAST2 (Bouckaert et al., 2014) on the non-polarized variants for all European isolates except WGRS 5, three Kenyan isolates (WGRS 26, WGRS 29 and WGRS-Test 23) and the American reference genome. Sample WGRS 5 was labeled as sample from Kenya, but population structure analyses clustered it with the Big Clonal cluster. We therefore excluded it from analyses using geographic information. Each isolate was timed by its year of isolation. As site model, we used a Γ model with four categories and estimated the proportion of invariant site, starting with a proportion of 0.8. We used the HKY model for mutation, estimating the frequencies and assumed used a strict molecular clock. Test runs with a relaxed exponential molecular clock with two discrete rates and with the different mutation model GTR showed only very small changes, which indicated that potentially shorter generation times of *S. turcica* in warmer climates need not to be accounted for. Since only in-species samples are included, we used the Coalescent Extended Bayesian Skyline as the tree model. As starting tree, we used the cluster tree estimated via `NJ2`. All other model settings were kept at the default values. The MCMC parameters were 225 million cycles (every 100th trace and 1,000th tree stored) with a 10 million pre-burnin period. For tree annotation, we used a 10 % burn-in. We conducted two independent runs. Effective Sample Size (ESS) scores, which estimate the numbers of independent draws from the posterior distribution that each BEAST run represents, were >100 for all non-population size parameters. Several population size parameter scored between 45 and 100 for a single run, all but one (=93) scored > 100 when runs were combined.

### Variation in sequence coverage

To use variation in sequence coverage as phylogenetic signal, we calculated variation of coverage in 1kb windows in each isolate. The most variable 2.5% windows were used to calculate a pairwise distance matrix of variance in coverage between all samples, from which a Neighbor-Joining tree was constructed.

### Reference-free association mapping

To characterize sequence reads that did not map to the reference, we conducted reference-free association study based on *k*-mers using the HAWK (Hitting Associations With *k*-mers) pipeline (Rahman et al.,2018). We ran HAWK between pairs of different clusters and between pairs of races independent of their assignment to populations: race 1 vs. race 0, race 0 vs. all, race 1 vs. all, race 3 vs. all, race 3N vs. race 0, race 3N vs. all, race 3 vs. race 1 and 0, race 3N vs. race 1 and 0, race 3 and 3N vs race 0 and 1. Race 1 vs. race 0 was also analysed for samples from the Big Clonal cluster only. Significantly differentiated *k*-mers were mapped against the reference genome to characterize the extent of clustering in some regions. Repetitive elements in windows with high number of *k*-mers mapped were searched with the protein-based RepeatMasking (Smit and Green, 2015). Gene-rich or gene-poor regions were determined for windows with high numbers of mapped *k*-mers by counting genes in these regions. Remaining unmapped assembled *k*-mers were compared against the NCBI non-redundant protein database using BLASTX to identify putative protein sequences.

## Data Availability

Raw sequence data of the 121 isolates generated in this study is available in the European Nucleotide Archive (ENA) under the project ID PRJEB37432. Scripts for analysing the data are availble from DOI: 10.5281/zenodo.4036236. Geographic and phenotypic information of the isolates is in the Supplementary Dataset S1.

## Acknowledgments

We thank Elisabeth Kokai-Kota for cultivating pathogen isolates, DNA extraction and sequencing preparation. We are grateful to Ana Galiano and Thomas Miedaner for comments on the manuscript. This work was funded by Deutsche Forschungsgemeinschaft (DFG) Priority program SPP1819 Rapid Evolutionary Adaptation (SCHM1354/11-1) to K. S. and SPP1590 Probabilistic Structures in Evolution (FR3633/2-1) to F. F. The authors acknowledge the additional support by the state of Baden-Württemberg through bwHPC.

## Author Contributions

FF, AvT and KS designed the study. AvT contributed materials. HH phenotyped the races. MV-V, FF, and KS analysed the data. MV-V, FF and KS wrote the manuscript. The authors declare no conflict of interest.

## Notes

http://dx.doi.org/10.5281/zenodo.4036236

### Competing Interest Statement

The authors have declared no competing interest.

### Summary of Updates

Revision includes only editorial changes on the manuscript. Title and abstract updated to better express the key message of the paper. Manuscript was revised to be more concise. No changes in the scientific content of the paper.

