## Supplemental Material for "Population genomic evidence for a repeated introduction and rapid expansion in Europe of a maize fungal pathogen"

### Supplementary text

#### (A) Simulation of Tajima's $D$ for Big Clonal and outlier test

For the outlier values of Tajima's $D$ for the clonal lineage Big Clonal, we compare the extreme positive values of Tajima's $D$ in 100 kb windows (Table 2) with simulations under the standard (Kingman) coalescent model for neutral evolution in an exponentially growing population. We consider growth rates $\{0.5, 5, 12.5, 25, 50, 500\}$ on a scale where $2N$ generations correspond to one unit of coalescent time.
We then perform 75,000 fixed-$s$ coalescent simulations given the number of segregating sites of the two windows (sample size 47) using the `phyclust` package in `R`. Then, we perform a Monte Carlo test for a positive outlier, i.e. the $p$-value is given by $\frac{1+H}{1+\#sims}$, where $H$ is the number of simulations where Tajima's $D$ was at least the value observed in the window. Unadjusted $p$-values are for the outliers in scaffolds 4 and 10, and after Bonferroni-Holm correction for multiple testing (when performing tests for all 340 windows of size 100 kb on the 15 biggest scaffolds). Here, the Bonferroni-Holm correction is conservative since we will have positive correlation between Tajima's $D$ values in nearby windows.

#### (B) Additional BEAST runs

We performed two additional BEAST analyses to test for the amount of bias introduced by including recombining clusters. For the first additional analysis (analysis I), we included only the Big Clonal (excluding WGRS_5), Small Clonal, and French Clonal clusters and the reference genome in a BEAST analysis. The Kenyan and Diverse clusters were excluded from the analysis to omit most recombination signals that were not incorporated in the BEAST approach, following Latorre et al. (2020). Apart from samples used and prolonged MCMC runs (300 million iterations), we performed analysis I analogously to the BEAST analysis described in the main manuscript. The second additional analysis (analysis II) used non-prolonged MMC runs (225 million iterations), a reduced burn-in of 5 million iterations and further omitted the reference genome from the used samples in comparison. For both BEAST runs of analysis I, ESS scores were >200 for all but population size parameters, of which three were <200 (minimum ESS of 172). For both BEAST runs of analysis II, ESS scores were below 100 for one and ¡200 for several other population size parameters, but above 200 for all other parameters. Analysis I obtained the same phylogeny as the main BEAST analysis (Fig. 4A, main manuscript) and split time estimates that are slightly more in the past (Table 4, Figs. 9, 10). Analysis II showed much noisier estimates than Analysis I (Table 4), in contrast to similar BEAST setups in Latorre et al. (2020) for *Magnaporthe oryzae* and despite using samples with fewer recombination events between one another. A potential reason for this is that without the reference and/or further sequences, the narrow time window of sampling and the low within-lineage segregation do not allow to date the phylogeny well. Moreover, compared to the analysis of the phylogenetic relationships between *M. oryza* individuals from Latorre et al. (2020), our phylogenetic network shows fewer need to add reticulate events to explain the genetic diversity, thus the bias introduced by non-modelled reticulation is likely smaller. We thus discarded the analysis II from further interpretation. The similar results of analysis I and the BEAST analysis from the main manuscript show that the results from the main analysis are not biased meaningfully by the inclusion of further recombination events and thus are robust with respect to the mode of reproduction of *S. turcica*.

#### (C) Posterior predictive checks

To assess the goodness-of-fit of the exponential growth model for the Kenyan and Big clonal population, we simulated the model for each scaffold 75,000 times. As summary statistics to check the fit, we used nucleotide diversity, Tajima's $D$ and the folded site frequency spectrum $fS_1, \ldots, fS_K$. For the latter, we record the number of SNPs $fS_i$ with minor allele count $i$ and $fS_K+$ as the number of SNPs with allele count $\geq K$ and set $K$ to the minimum of 15 and the possible maximum minor allele count in the sample. The growth rate was set to the estimated posterior mean growth rate from Table 3 (main manuscript)and mutation rate set to the generalized Watterson estimate based on the number of SNPs in the scaffold. Then, the actually observed statistics are compared with the distribution of values simulated from the fitted model, see 12 and 11.

#### (D) ABC model selection for exponentially growing populations: Multiple merger coalescents vs. Kingman's coalescent

Following the approach from Menardo et al. (2020), we perform additional model selection between Kingman's $n$-coalescent with exponential growth and Beta-$n$-coalescents and Dirac-$n$-coalescents with exponential growth. We use the same prior and parameter range for Kingman's $n$-coalescent as in the ABC analysis from the main manuscript. For coalescent parameter ranges and priors for the multiple merger coalescents with growth, we take discrete uniform priors for the coalescent parameters $p \in \{0.05, 0.15, \ldots, 0.95\}$ and $\alpha \in \{1, 1.1, \ldots, 1.9\}$, additionally the growth parameter has a discrete $\log$-uniform distribution on $[0, 50]$ with 10 steps $\{0, 1.54, 2.39, \ldots, 20.9632.37, 50.00\}$ (rounded to two digits). Simulations are performed as described in (Menardo et al., 2020), the ABC model selection is performed as described in the methods section of the main manuscript. The range of mean out-of-bag errors was comparable to the model selection in the main text with 23-24% for Big Clonal and 19-20% for the Kenyan cluster. Due to low posterior probabilities (Big Clonal: 51%-61%, Kenyan: 58%-66%) and thus inconclusive evidence for which model fits better (Big Clonal: 3 scaffolds support Beta coalescent with growth, 2 scaffolds support Kingman's coalescent with growth; Kenyan: all 5 scaffolds support Kingman's coalescent with growth), we did neither perform posterior predictive checks nor estimate parameters for multiple merger models with growth.

#### (E) Sharing the MRCA between sample and population

Following (Saunders et al., 1984, Eq. 4.5), under a neutral (standard) coalescent setting with arbitrary demographic changes (which lead only to a time-change of the tree), the probability that the most recent common ancestor (MRCA) of a sample of size $n$ does not change regardless of how many more individuals are sampled, is $\frac{n-1}{n+1}$. Applied to the three clonal lineages, this leads to probabilities of (Big Clonal $\approx 0.96$ , Small Clonal $\approx 0.88$, French Clonal $\approx 0.82$) that the estimated times of emergence based on a sample reflect the time back to the MRCA of the lineages.

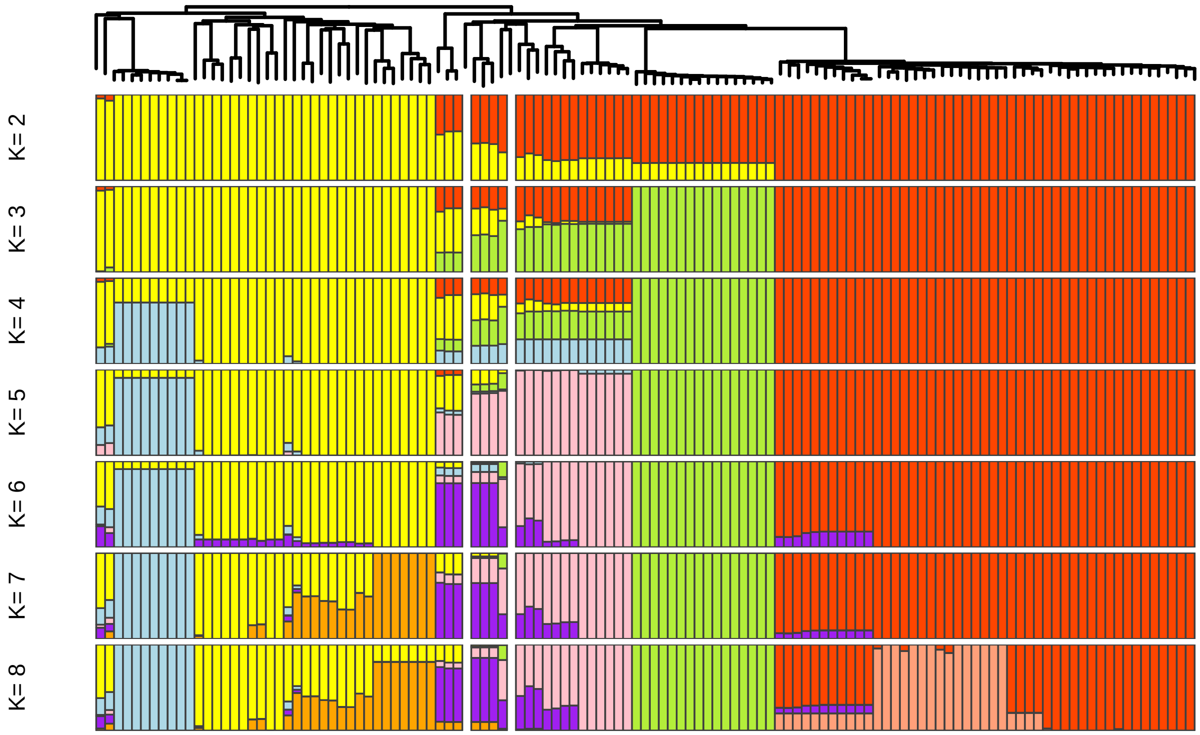


Figure 1: Individual ancestry coefficients for different K values in the same order as the rooted NJ tree.

Table 1: Differences between technical replicates as the number of variants between them

| isolate | # SNP replicates | total SNP dataset | % SNP replicates |
|---|---|---|---|
| WGRS_0586 | 7 | 23209 | 0.030 |
| WGRS_1200 | 28 | 23209 | 0.121 |
| WGRS_1205 | 242 | 23209 | 1.043 |
| WGRS_1231 | 6 | 23209 | 0.026 |
| WGRS_1235 | 16 | 23209 | 0.069 |
| WGRS_1243 | 12 | 23209 | 0.052 |
| WGRS_1263 | 7 | 23209 | 0.030 |
| WGRS_1480 | 4 | 23209 | 0.017 |

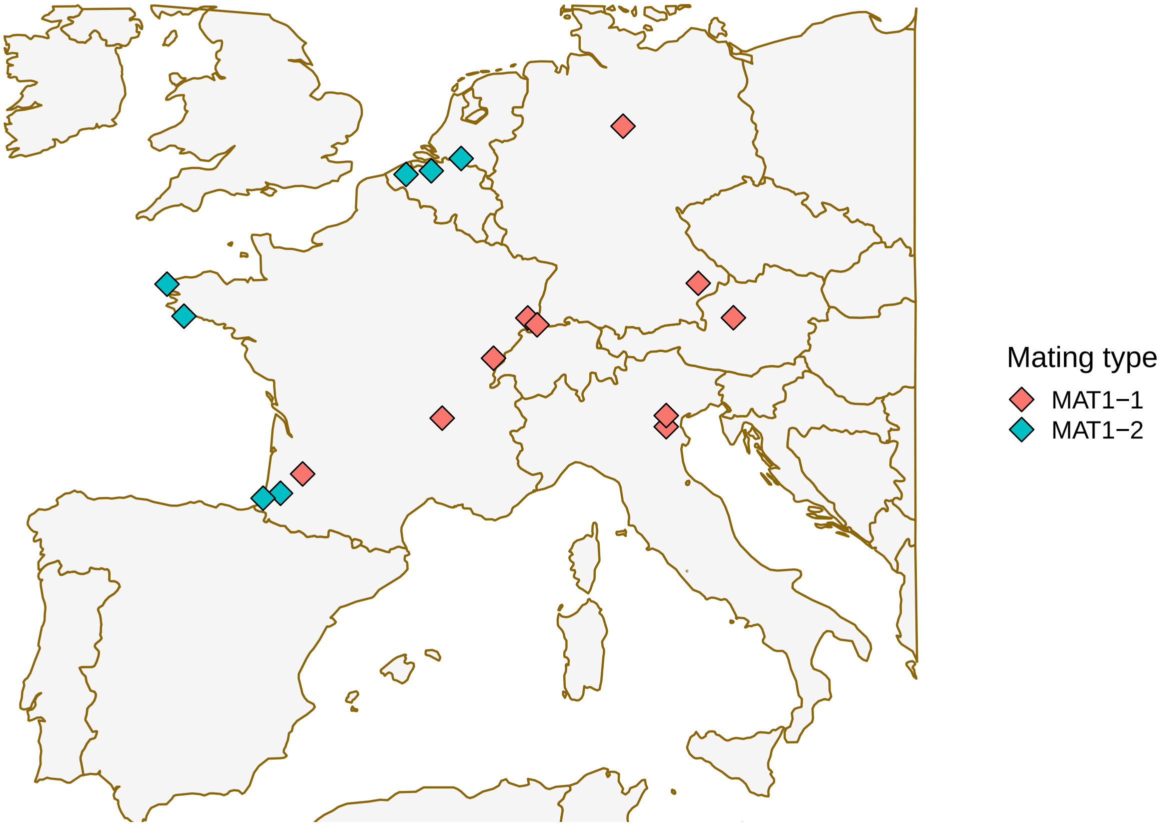


Figure 2: Geographic distribution of the mating type in the Diverse cluster.

Table 2: Tajima's $D$ outlier windows for Big Clonal and Small Clonal - diversity statistics

| Pop | Scaffold | Start | $\pi$ | Qu | $\theta_w$ | Qu | $D$ | Qu | $HD$ | Qu |
|---|---|---|---|---|---|---|---|---|---|---|
| Big Clonal | 4 | 1600001 | 0.44 | 0.83 | 0.23 | 0.28 | 1.31 | 1.00 | 0.44 | 0.86 |
| Big Clonal | 10 | 500001 | 0.90 | 0.95 | 0.45 | 0.51 | 1.80 | 1.00 | 0.51 | 0.92 |
| Small Clonal | 7 | 1700001 | 0.46 | 0.92 | 0.30 | 0.81 | 1.03 | 0.99 | 0.46 | 0.94 |
| Small Clonal | 10 | 1000001 | 0.52 | 0.95 | 0.30 | 0.81 | 1.47 | 1.00 | 0.52 | 0.95 |
| Small Clonal | 15 | 800001 | 0.46 | 0.92 | 0.30 | 0.81 | 1.03 | 0.99 | 0.46 | 0.94 |

All windows (100 kb) with most elevated D across the 15 biggest scaffolds are shown per population ($D \geq 1$). Qu: Quantile of window diversity (statistic on the left) among all windows in Scaffolds 1-15. $\pi$: nucleotide diversity, $\theta_w$: Watterson's estimate, HD: haplotype diversity.

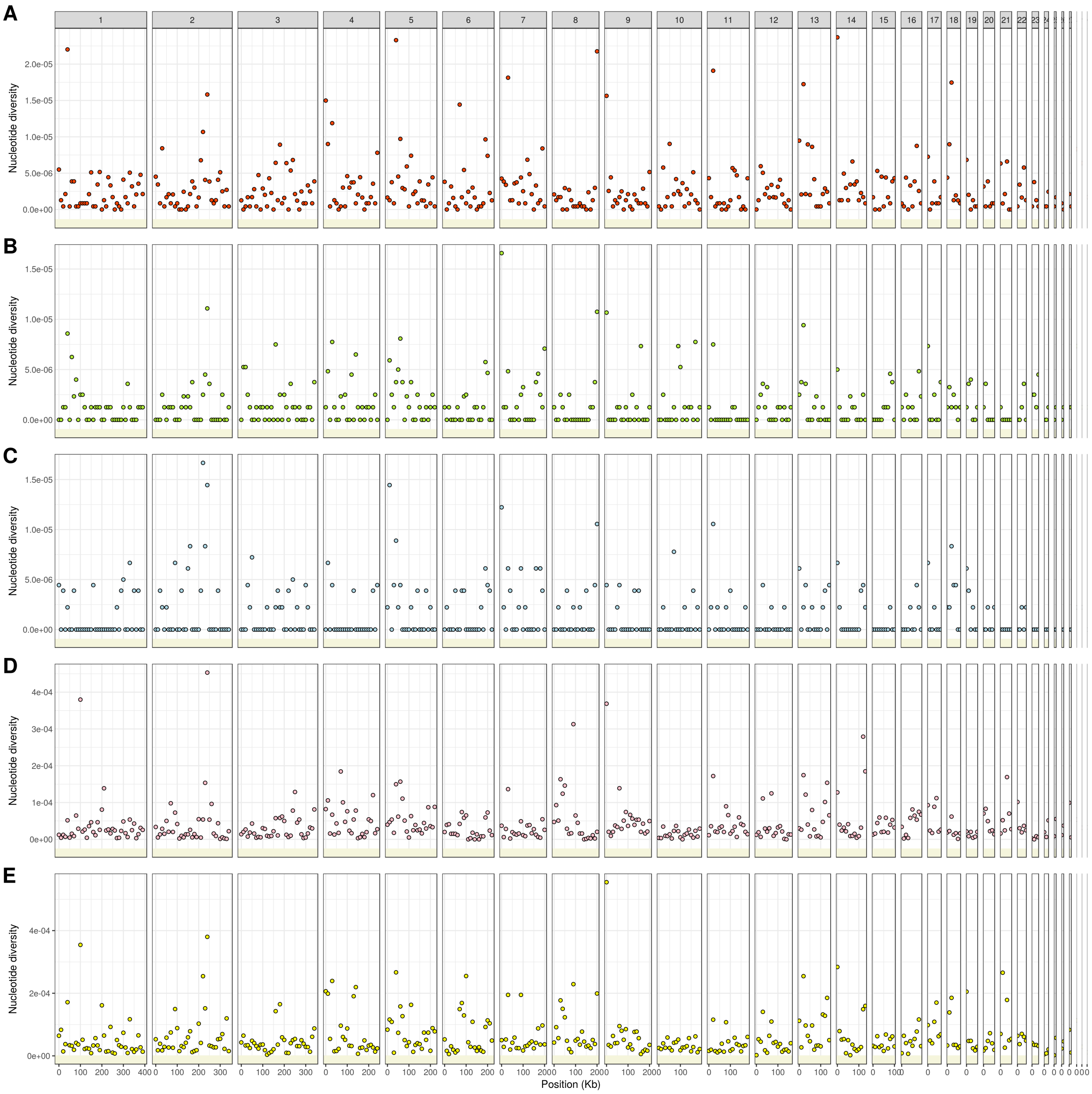

Figure 3: Nucleotide diversity per bp across the biggest 15 scaffolds (ordered by size) in 100k bp windows for **A**) Big Clonal, **B**) Small Clonal, **C**) French Clonal, **D**) Diverse and **E**) Kenyan. Negative values in yellow area indicate missing data.

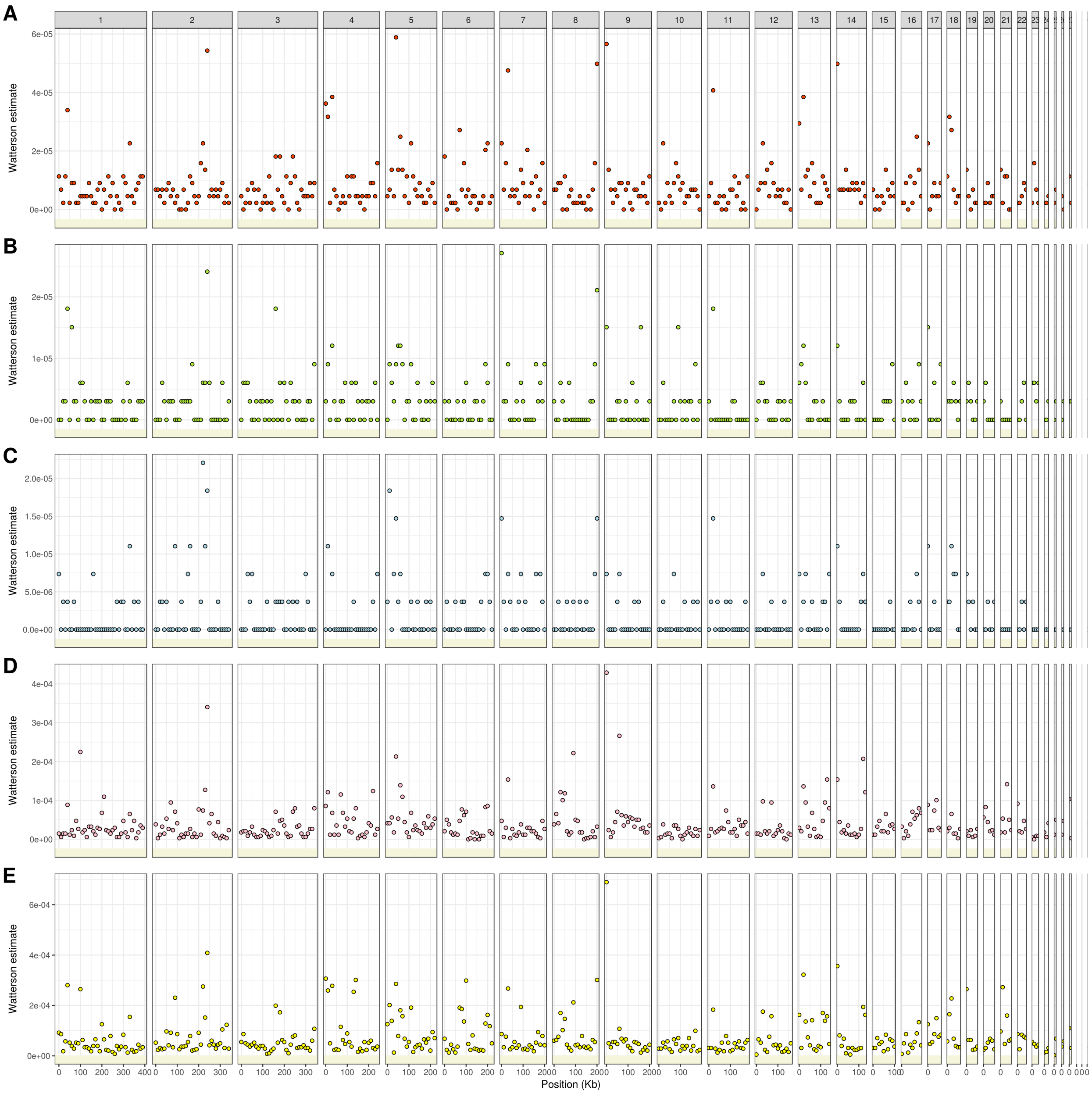


Figure 4: Watterson's estimator per bp across the biggest 15 scaffolds (ordered by size) in 100k bp windows for **A**) Big Clonal, **B**) Small Clonal, **C**) French Clonal, **D**) Diverse and **E**) Kenyan. Negative values in yellow area indicate missing data.

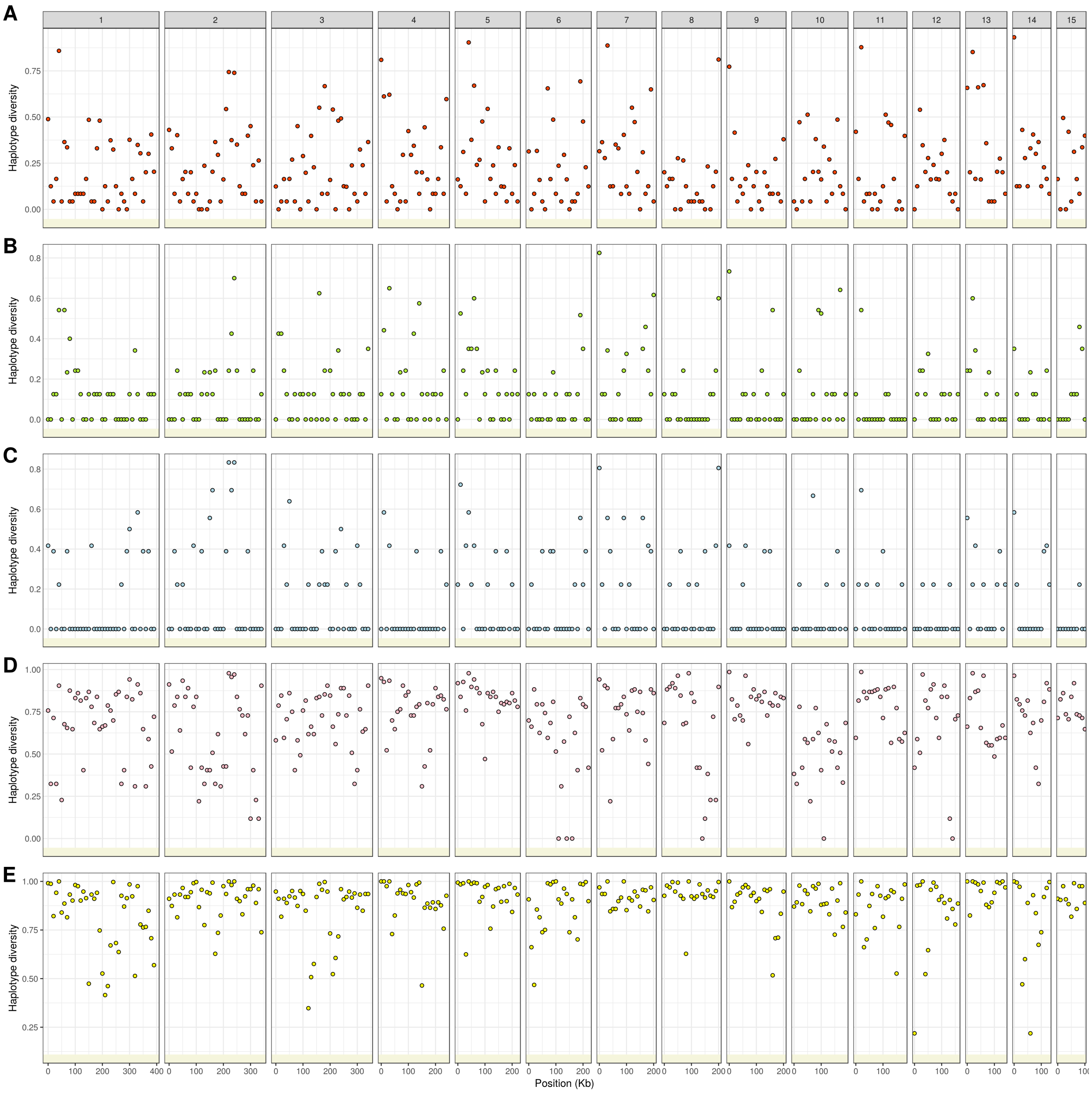


Figure 5: Haplotype diversity across the biggest 15 scaffolds (ordered by size) in 100k bp windows for **A**) Big Clonal, **B**) Small Clonal, **C**) French Clonal, **D**) Diverse and **E**) Kenyan. Negative values in yellow area indicate missing data.

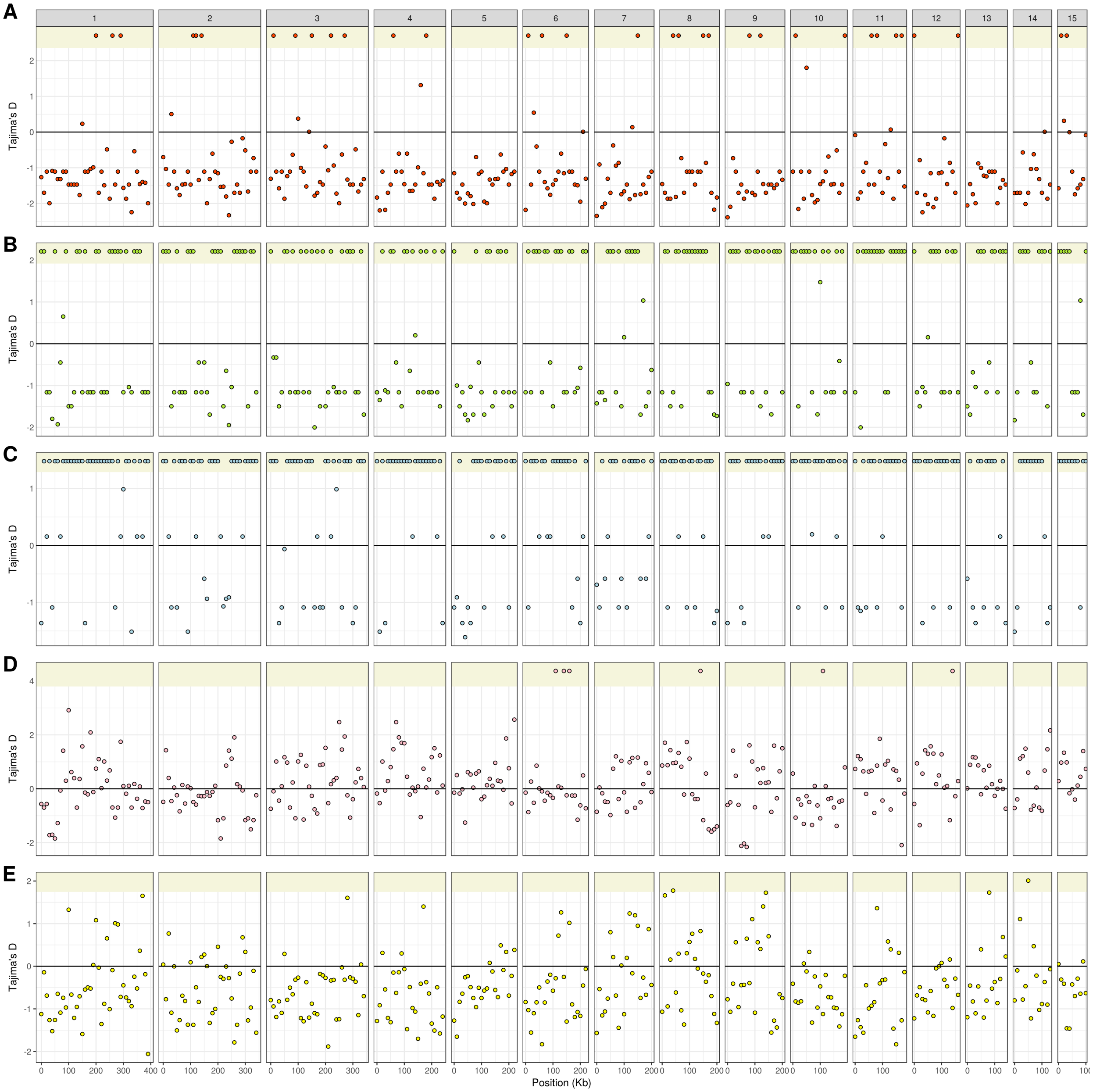


Figure 6: Tajima's $D$ across the biggest 15 scaffolds (ordered by size) in 100k bp windows for **A**) Big Clonal, **B**) Small Clonal, **C**) French Clonal, **D**) Diverse and **E**) Kenyan. Windows in the yellow area above 2 indicate missing data.

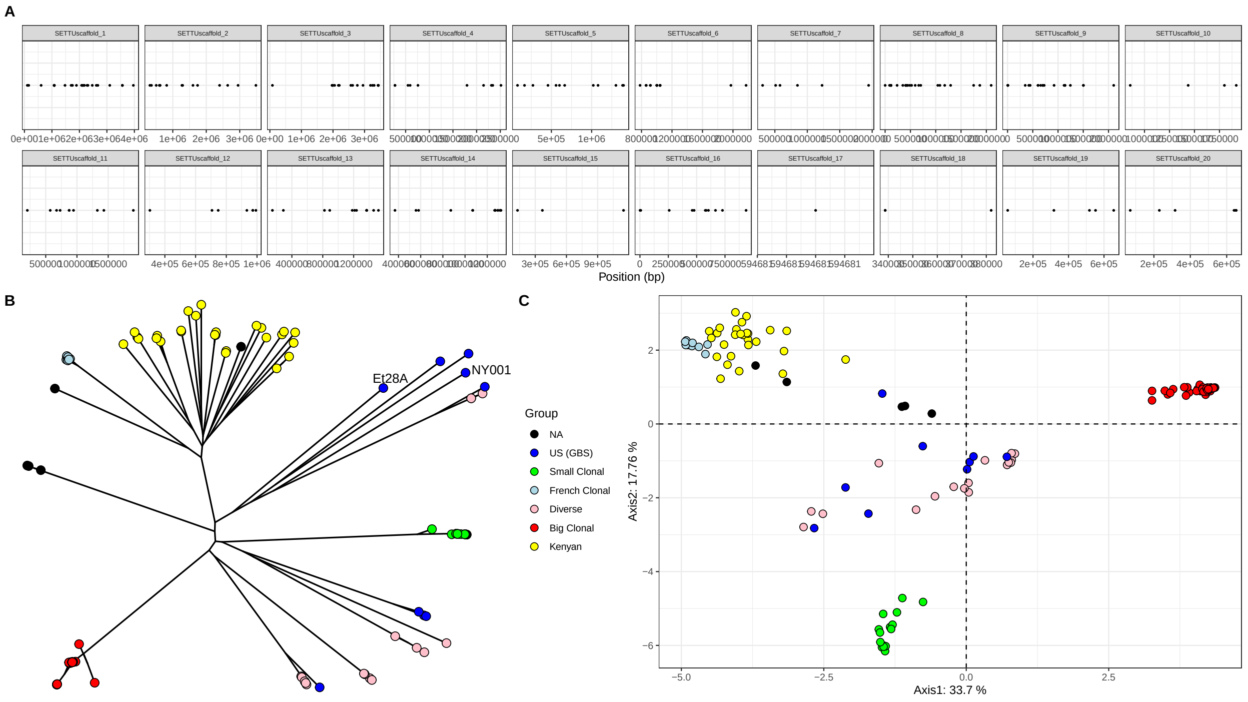


Figure 7: **A**) SNP distribution (first 20 scaffolds) **B**) NJ Tree and **C**) PCA of the 280 SNPs from the merged dataset with GBS samples from US

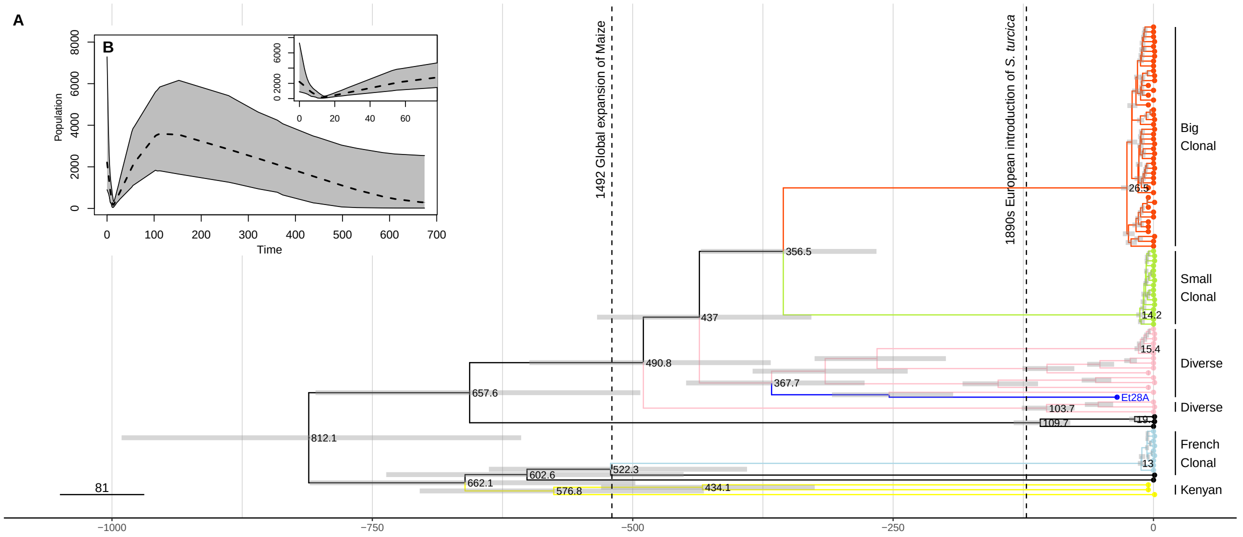


Figure 8: **Beast replication**. **A**) Dated phylogeny obtained with BEAST, using all European isolates, reference genome Et218A (in dark blue) and three samples from Kenyan cluster. Time is given as years before 2012, the year of the most recent sampling. Horizontal gray bars show the 95% highest posterior density intervals (95% HPI) for split times. **B**) Extended Bayesian skyline plot obtained with BEAST for the analysis from (A). The inset zooms into the most recent past. Time runs backwards from 2012. The dashed line shows the median posterior population size, while the gray area shows the 95% HPI.

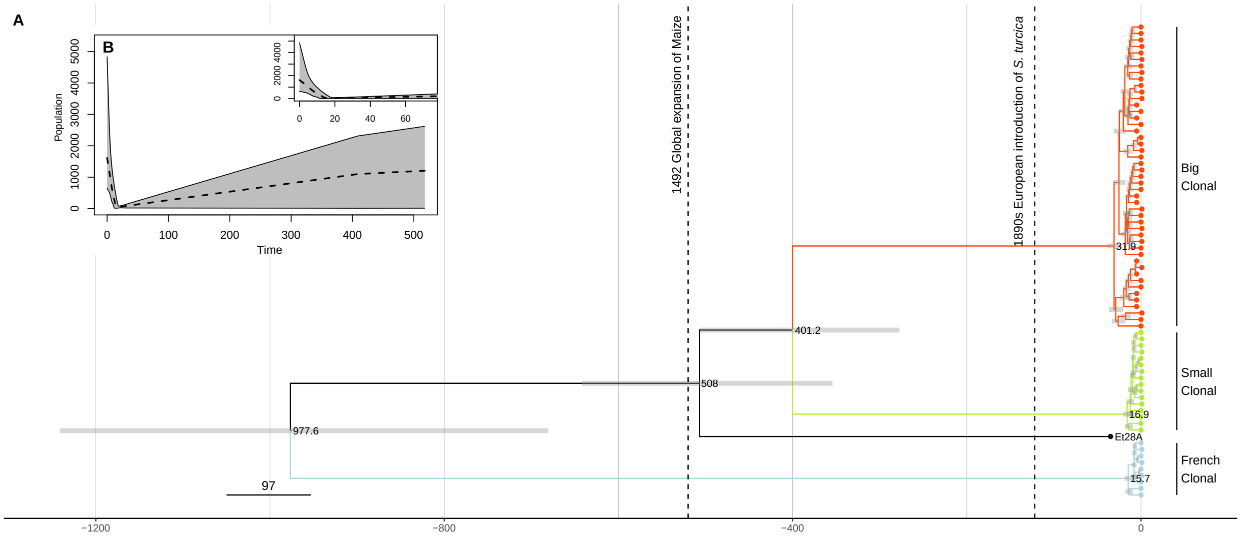


Figure 9: **A**) Dated phylogeny obtained with BEAST, using all isolates from the three European clonal lineages Big Clonal, Small Clonal, French Clonal and the genome reference (Et28A). Time is given as years before 2012, the year of the most recent sampling. Horizontal gray bars show the 95 % highest posterior density intervals (95% HPI) for split times. **B**) Extended Bayesian skyline plot obtained with BEAST for the analysis from (A). The inset zooms into the most recent past. Time runs backwards from 2012. The dashed line shows the median posterior population size, while the gray area shows the 95% HPI.

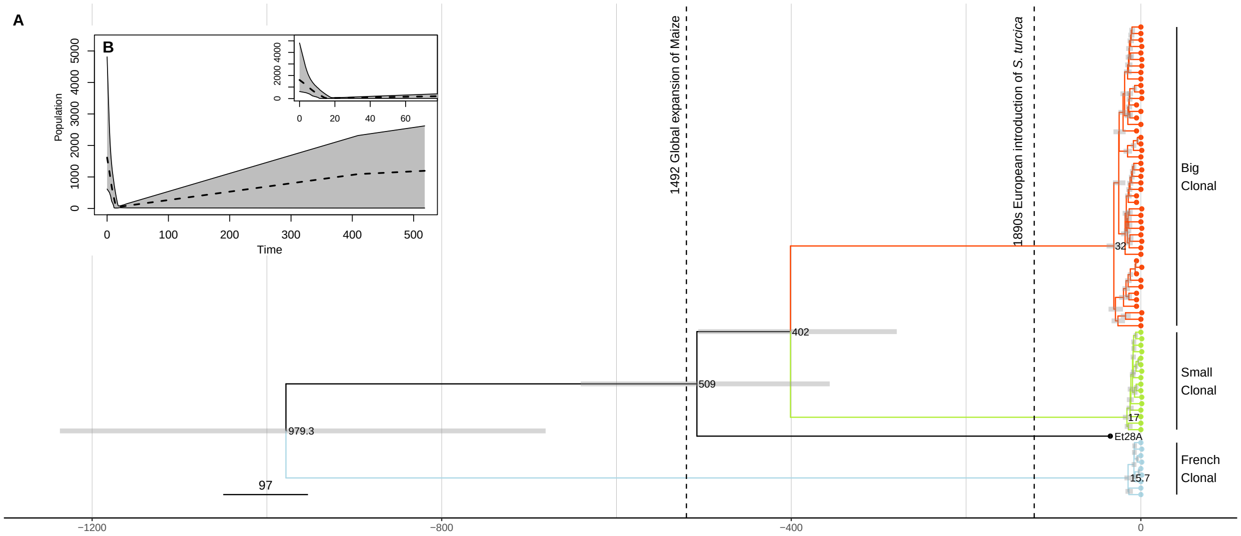


Figure 10: **A**) Dated phylogeny obtained with BEAST (replication run), using all isolates from the three European clonal lineages Big Clonal, Small Clonal, French Clonal and the genome reference (Et28A). Time is given as years before 2012, the year of the most recent sampling. Horizontal gray bars show the 95 % highest posterior density intervals (95% HPI) for split times. **B**) Extended Bayesian skyline plot obtained with BEAST for the analysis from (A). The inset zooms into the most recent past. Time runs backwards from 2012. The dashed line shows the median posterior population size, while the gray area shows the 95% HPI.

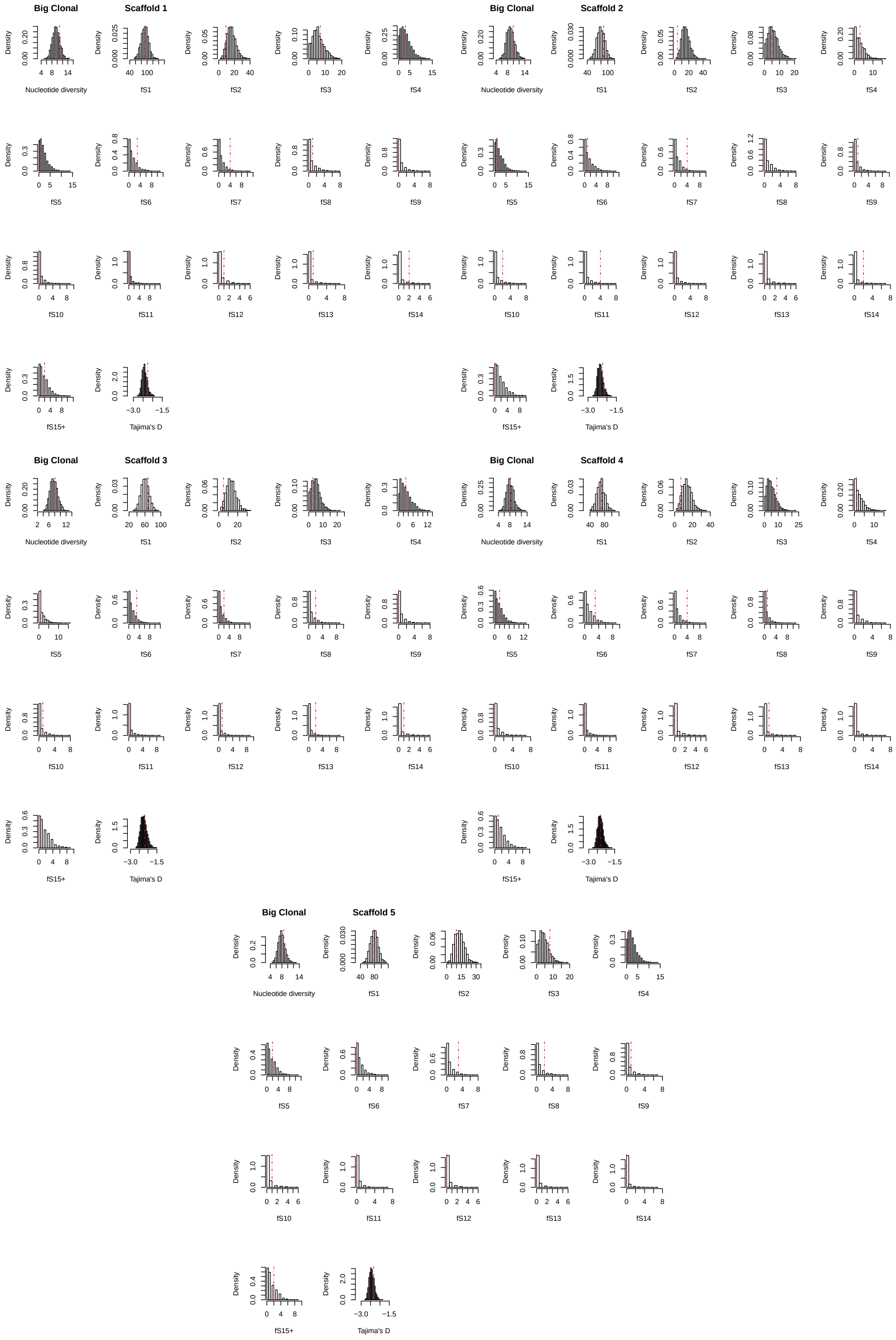


Figure 11: Posterior predictive checks under the fitted exponential growth model for lineage Big Clonal

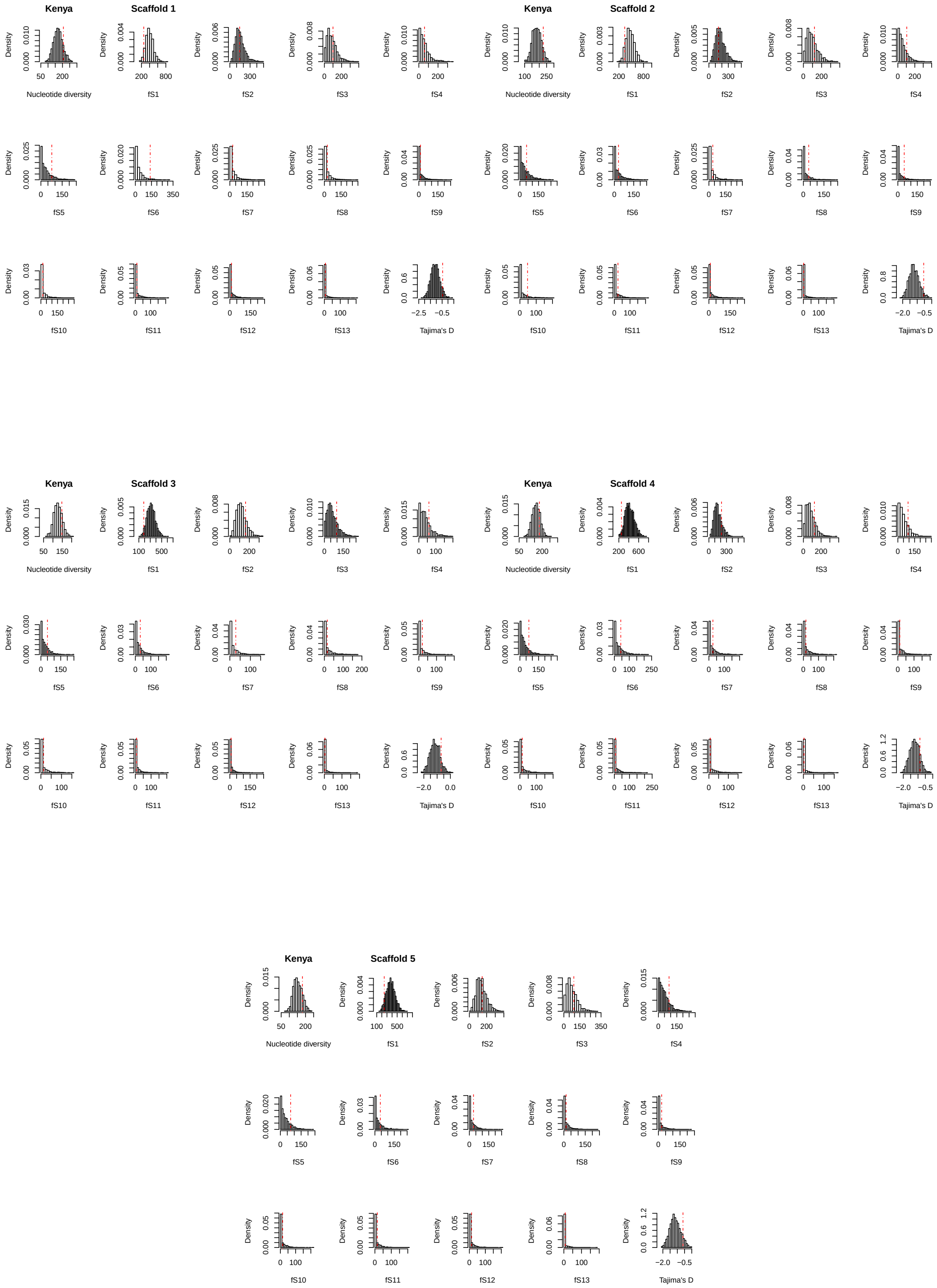


Figure 12: Posterior predictive checks under the fitted exponential growth model for the Kenyan population

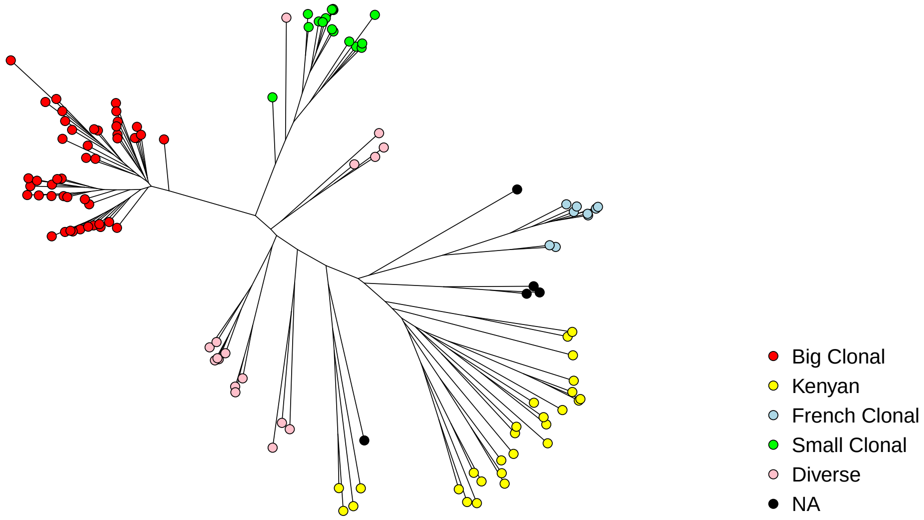


Figure 13: NJ Tree based on the sequence coverage of genomic regions to indicate the contribution of structural variants to genetic differentiation. Color according to defined clusters

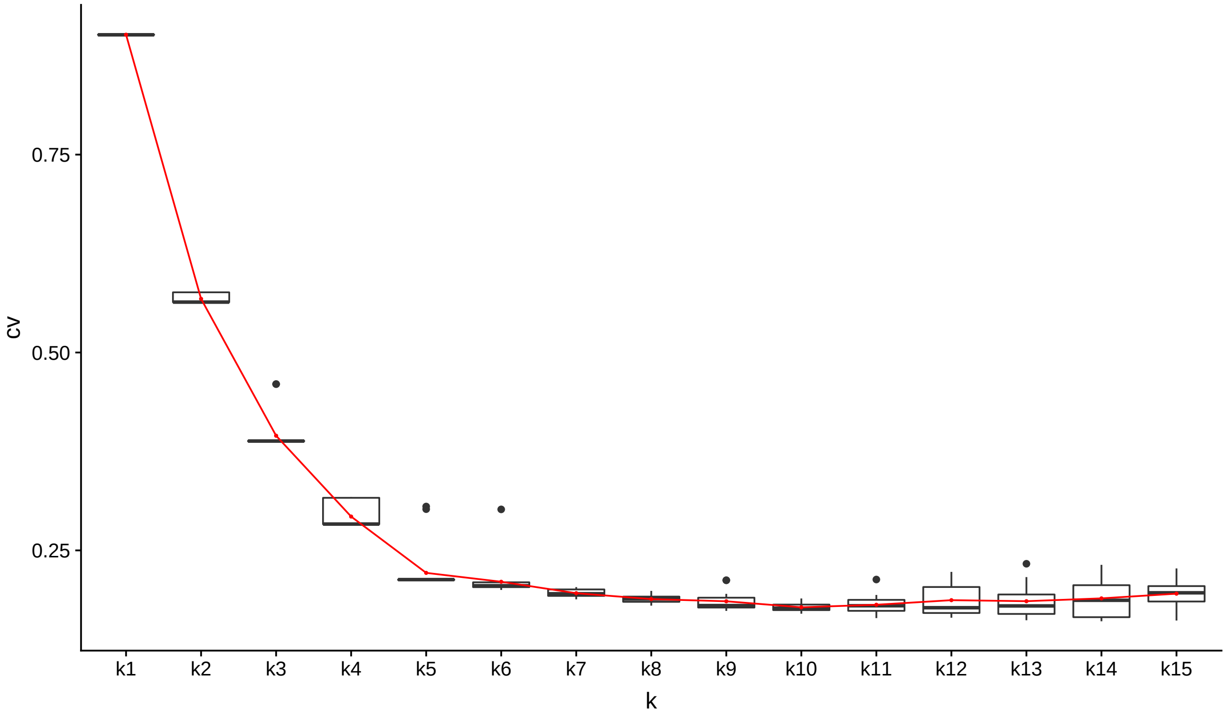


Figure 14: Admixture cross-validation error on 20 runs and up to 15 clusters.

Table 3: Tajima's $D$ outlier in Big Clonal - deviation from fitted exponential growth model

| Outlier scaffold | $g = 0.5$ | $g = 5$ | $g = 12.5$ | $g = 25$ | $g = 50$ | $g = 500$ |
|---|---|---|---|---|---|---|
| 4 | 0.143 (1) | 0.079 (1) | 0.053 (1) | 0.039 (1) | 0.030 (1) | 0.014 (1) |
| 10 | 0.035 (1) | 0.010 (1) | 0.005 (1) | 0.003 (0.884) | 0.001 ( 0.431) | 0.000 (0.073) |

$p$-values for Tajima's $D$ outliers in population Big Clonal are computed for comparison with models with exponential growth
(rate $g$) as described in Sect. . In brackets: $p$-value after correction for multiple testing. Values are rounded to three digits.

Table 4: Split and times to most recent common ancestor estimated with BEAST

| Split/MRCA | (BC,SC,FC,REF) | (BC,SC,FC) | (BC,SC,FC,REF,D,Kenyan) |
|---|---|---|---|
| Big Clonal (BC) | 31.93 [24.21,41.92] | 71.62 [37.75,145.8] | 26.6 [21.21,33.44] |
| Small Clonal (SC) | 16.95 [12.58,22.58] | 39.05 [20.29,79.74] | 14.23 [10.95,18.38] |
| French Clonal (FC) | 15.67 [11.79,20.83] | 34.56 [17.73,71.03] | 13.03 [10.13,16.63] |
| Diverse (D) | - | - | 492.05 [388.47,625.83] |
| (BC,SC) | 401.61 [303.74,531.77] | 1059.41 [547.07,2171.59] | 357.43 [282.44,454.8] |
| (REF,(BC,SC)) | 508.53 [383.04,670.62] | - | 438.2 [290.92,466.81] |
| (D,(BC,SC)) | - | - | 438.19 [345.09,555.08] |
| (FC,Kenyan) | - | - | 663.83 [522.57,844.5] |
| (FC,(BC,SC,REF)) | 978.43 [731.67,1294.4] | 2200.24 [1143.21,4518.04]* | |
| ((FC,Kenyan),(BC,SC,D)) | - | - | 814.35 [644.33,1036.73] |

In each cell, the first number shows posterior median, in parentheses 95% HCI. Times are measured in years back from the latest sampling date (2012).The first column shows the same for the analysis based on the samples from the three clonal lineages and the reference genome. The third column shows the mean median split/emergence times from two runs with the 'full setup' (all but one European samples, 2 Kenyan and the reference) and the union of the 95% HCIs for these two BEAST runs. The second column shows the result for a single BEAST run when using only the data from the clonal lineages. $^{*}$: corresponds to the estimated split time between (Big Clonal ,Small Clonal) and French Clonal.

Table 5: RF-ABC model selection results

| Population | SNP count | Mean OOB | Best model | Post. prob. best model | Fitted parameter |
|---|---|---|---|---|---|
| Small Clonal | 24-34 | 34-36% | 3x EXP | 82-95% | $g = 115$-1189 |
| | | | 2x Dirac | 63-68% | $p = 0.53$-0.59 |
| French Clonal | 9-28 | 39-46% | 5x EXP | 32 - 90% | $g = 33$-209 |
| Diverse | 301-466 | 25-26% | 4x EXP | 50 - 57 % | $g = 0.7$-1 |
| | | | 1x Beta | 51 % | $\alpha = 1.66$ |

The table shows ABC results for the five largest scaffolds in different lineages. Mean OOB is the out-of-bag prior error rate for model classes, averaged over all model classes. Best model summarises the counts of best fitting models across the 5 scaffolds (EXP: Kingman's $n$-coalescent with exponential growth, Dirac: Dirac-$n$-coalescent, Beta: Beta-$n$-coalescent). Posterior prob. best model gives the posteror probability of the best fitting model(s). Fitted parameter: $g$ is the posterior median of the exponential growth parameter in coalescent units, where one unit represents $2N$ generations, $p$ the coalescent parameter of the Dirac $n$-coalescent and $\alpha$ the coalescent parameter of the Beta-$n$-coalescent. For each variable, we report the range of values for the five biggest scaffolds.

Table 6: $k$-mer analysis with HAWK on different group of races

| Races case | Races control | # samples case | # samples control | # $k$-mer (assembled) case | # $k$-mer (assembled) control | # BLAST hits |
|---|---|---|---|---|---|---|
| 1 | 0 | 13 | 30 | 0 | 168 (3) | 1 ** |
| 3 | 0 | 8 | 30 | 0 | 0 | - |
| 3N | 0 | 6 | 30 | 0 | 0 | - |
| 1 * | 0 * | 9 | 20 | 0 | 0 | - |
| 0 | 1, 3, and 3N | 30 | 27 | 10 (0) | 0 | 0 |
| 1 | 0, 3, and 3N | 13 | 44 | 0 | 0 | - |
| 3 | 0, 1, and 3N | 8 | 43 | 0 | 0 | - |
| 3N | 0, 1, and 3 | 6 | 51 | 0 | 0 | - |
| 3 | 0 and 1 | 8 | 43 | 0 | 0 | - |
| 3N | 0 and 1 | 6 | 43 | 0 | 0 | - |
| 3 and 3N | 0, Race 1 | 14 | 43 | 0 | 12 (0) | 0 |

* Samples only from Big Clonal cluster. ** Hit name: polyketide systase protein

#### Supplementary Figures

#### Supplementary Tables
